# Pax6 and KDM5C co-occupy a subset of developmentally critical genes including Notch signaling regulators in neural progenitors

**DOI:** 10.1101/579219

**Authors:** Giulia Gaudenzi, Olga Dethlefsen, Julian Walfridsson, Ola Hermanson

## Abstract

Pax6 is a key transcription factor in neural development. While generally viewed as a transcriptional activator, mechanisms underlying Pax6 function as a repressor is less well understood. Here we show that Pax6 acts as a direct repressor of transcription associated with a decrease in H3K4me3 levels. Genome wide analysis of the co-occupancy of the H3K4 demethylase KDM5C and Pax6 with H3K4me3-negative regions revealed 177 peaks on 131 genes. Specific analysis of these Pax6/KDM5C/H3K4me3-peaks unveiled a number of genes associated with Notch signaling, including *Dll1, Dll4*, and *Hes1*. RNA knockdown of PAX6/KDM5C in human neural progenitors resulted in increased *DLL4* gene expression, decreased *DLL1* expression, and no significant effect on HES1 mRNA levels, differences that could be due to gene-specific variations in the chromatin landscape. Our findings suggest that PAX6 and KDM5C co-regulate a subset of genes implicated in brain development, including members of the Notch signaling family.

---

The proper development of the mammalian cerebral cortex is strictly dependent on precise spatial and temporal intracellular and extracellular cues^1,2^. Genetic studies of psychiatric disorders have revealed that many of the factors mutated in such disease are regulators of neural development^3^. Further, a subset of these genes harboring critical mutations has been shown to be regulators of chromatin, i.e. epigenetic regulators^4^. Indeed, genetic defects in epigenetic regulators seem to be common in neurodevelopmental disorders such as Rett syndrome, Rubinstein-Taybi syndrome, and other states of mental retardation^5,6^.

It has further been shown that valproic acid (VPA), but not lithium, are teratogenic, and offspring of mothers taking VPA show higher frequency of autistic spectrum disease^7^. This is a highly relevant observation as VPA and lithium share many signaling mechanisms^8^ but while VPA is in addition an inhibitor of histone deacetylase (HDAC) activity, lithium is not^9,10^. Altogether, these reports highlight the importance of appropriate function of histone modifying proteins in neural development.

We and others have previously demonstrated specific roles for HDACs, histone acetyl transferases (HATs), associated co-activators and co-repressors, as well as DNA methylation regulators in many events during neural development, including differentiation, proliferation, cell death, senescence, and autophagy^11-17^. These factors have in addition been linked to DNA-binding transcription factors involved in neurodevelopmental disorders and fundamental neurodevelopment, such as REST and MeCP2^18-23^.

During the past years, roles for factors regulating histone methylation has emerged^24,25^. One such histone demethylase that has previously been linked to psychiatric disease, especially X-linked mental retardation, is KDM5C (also known as SMCX, JARID1C)^26^. KDM5C is a demethylase of lysine 4 on histone H3 (H3K4). H3K4 trimethylation (H3K4me3) is strongly and directly associated with transcriptional activation^27,28^ and KDM5C is therefore primarily regarded as an inhibitor of transcriptional activation. KDM5C has more recently been linked to other psychiatric disorders including autism, in line with its presumed general importance for proper brain development^29^. Jumonji domain-proteins similar to KDM5C interact with sequence-specific transcription factors^18^, but less is known regarding such KDM5C-recruiting factors.

PAX6 is a DNA-binding transcription factor that has been implicated in the fundamental regulation of the development of the cerebral cortex^30,31^. Mutations in PAX6 are most commonly associated with aberrations in eye development, but studies have demonstrated that mutations in PAX6 can also be associated with neurological and psychiatric conditions^32^. Although PAX6 has long been studied from a genetic point of view, relatively little is known of the mechanisms underlying its function. It has been shown to act as a transcriptional activator, and in this role associated with H3K4me3 methyl transferases^33^. However, there are further reports of PAX6 preferentially binding to methylated DNA^34^, and when Pax6 is ablated, the expression of a subset of genes has been shown to be increased^35^. Indeed, it has recently been reported that PAX6 can interact with class I HDACs, specifically HDAC1, suggesting that PAX6 may mediate direct repression of transcription^36^. The increased gene expression could however theoretically be explained by a lack of activation of a secondary repressor, the association with binding to methylated DNA could be mere association, and HDAC1 is not ubiquitously expressed in neural progenitors^11^. There is thus a need for increased understanding of putative mechanisms underlying Pax6 function as a transcriptional repressor.

The aim of this study was to analyze Pax6 occupancy patterns and compare these to H3K4me3 levels and binding of the H3K4 demethylase KDM5C. Combining chromatin immunoprecipitation (ChIP) assays with sequencing (ChIP-Seq) is a powerful method to identify genome-wide DNA binding sites for transcription factors and other proteins. To this end we have downloaded and analysed two published ChIP-seq data sets^37,38^. While we found a subset of Pax6 peaks coinciding with H3K4me3-positive regions, most Pax6 binding was detected in regions with low H3K4me3 (H3K4me3-). Indeed, transcriptional assays showed that PAX6 could act as a direct repressor of transcription, and this repression was associated with a small but reproducible decrease in H3K4me3 levels. Analysis of the occupancy of KDM5C at regions with high and low levels of H3K4me3 revealed that KDM5C occupied H3K4me3-positive loci at a majority of peaks, but as expected, a large number of the KDM5C-occupied regions were H3K4me3-. We therefore made a comparison of all three datasets and found 177 peaks on 131 genes where Pax6 and KDM5C overlapped with H3K4me3-regions. When analyzing the function of the Pax6/KDM5C-occupied, H3K4me3-genes, we found a number of genes associated with Notch signaling, including *Dll1, Dll4*, and *Hes1*. RNA knockdown of PAX6 and KDM5C in human iPS-derived neural progenitors (hNPs) resulted in an increase in DLL4 gene expression, whereas no significant effect of HES1 mRNA levels was detected, and DLL1 gene expression actually decreased. Our results suggest that PAX6 and KDM5C are required components for proper regulation of gene expression of Notch signaling factors.

## RESULTS

### Genome wide analysis reveals Pax6 occupancy on H3K4me3–positive and –negative regions

Pax6 has been linked to transcriptional activation and shown to be associated with H3K4 methyl transferases^33^ whereas other reports have suggested that Pax6 may be linked to transcriptional repression^34,36^. Indeed, knockdown of Pax6 in neural progenitors (NPs) yields both increased and decreased gene expression (ref. 35 and data not shown). To investigate putative mechanisms underlying Pax6 function, we initiated an *in silico* study of genome wide Pax6 occupancy to compare with the H3K4me3 landscape in the rodent developing cortex using stringent bioinformatics analysis to interpret the results (**Supplementary file 1**). This initial analysis revealed that 9,352 ChIP-Seq peaks were positive for Pax6 binding (**Figure 1a**) in E12.5 mouse forebrain tissue^38^. When analyzing the enrichment of peaks near genes using the Reactome pathway terms, the results were predominantly associated to biological processes of development, axon guidance, and Notch signaling (**Figure 1b**) (over represented p-value 4.01E-08). Notch receptor family, its ligands, and signaling intermediate factors are essential for correct control of proliferation and differentiation, as well as other cellular events, in neural and organism development^39,40^. We then compared the Pax6 ChIP-seq peaks to regions rich in H3K4me3, and this analysis unveiled 15,745 H3K4me3-positive peaks. A large proportion of those peaks was associated with cell cycle control (**Figure 1a,c**) (over represented p-value 4.16E-54). This is completely in accordance with H3K4me3 as an active mark of transcription as most cells in an E12.5 mouse forebrain are dividing.

**Figure 1:**
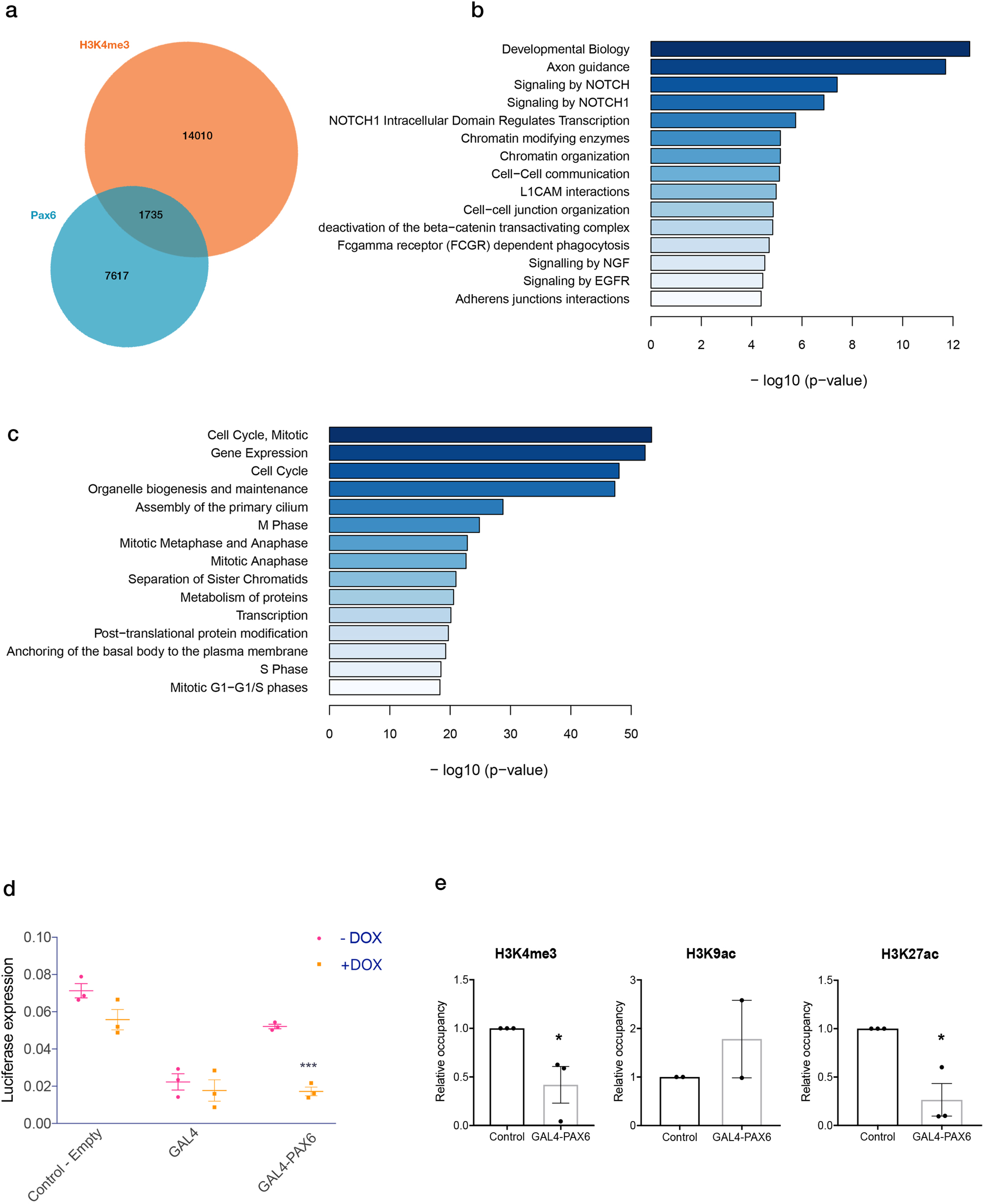
Pax6 occupancy is enriched at genomic regions with low H3K4me3 and Pax6 can exert repression associated with a decrease in H3K4me3 in transcriptional assays. (a) Genome wide analysis of ChIP-Seq experiments revealed that the vast majority of Pax6 peaks were found in regions with low H3K4me3. (b) Top Reactome terms for Pax6 peaks. (c) Top Reactome Terms for high H3K4me3 levels. (d) A Gal4-Pax6 construct exerts repression in a Dox-sensitive transcriptional assay. 2-way ANOVA shows the effect of DOX treatments (n (number of biological replicates) = 3 in each group. \*\*\**p*=0.0008, the data represent the mean ± SEM). (e) The repressor function of the Gal4-Pax6 fusion construct is associated with a decrease in H3K4me3 and H3K27ac but not H3K9ac levels.

When combining the two datasets, we found 1,735 overlapping peaks between Pax6 and H3K4me3 (**Figure 1a**). This confirms previous observations of Pax6 association with H3K4me3 methyl transferases^33^. The number of overlapping peaks was however lower than expected and the vast majority of Pax6-positive peaks were associated with H3K4me3-negative (H3K4me3-) regions (**Figure 1a, Supplementary file 1**).

### PAX6 can act as a repressor in transcriptional assays in association with decreased H3K4me3

The analysis of PAX6 occupancy in combination with H3K4me3-negative regions prompted us to investigate whether PAX6 could act directly as a repressor in transcriptional assays. We first constructed a Gal4-PAX6 fusion construct that could be used in a doxycycline (DOX) sensitive assay (DOX-on)^41^ (**Supplementary figure 1**), and analyzed its activation of a UAS-promoter in a luciferase reporter context in stably transfected HEK293 cells. This assay revealed that PAX6 indeed can act as a direct transcriptional repressor (**Figure 1d**) (*p*=0.0008), in accordance with recent observations^36^.

We next utilized the DOX-on system to investigate the effects on chromatin modifications on the stably transfected promoter (**Figure 1e**). We analyzed three of the most well-studied lysine modifications, i.e. H3K4me3, acetylation of lysine 9 on histone H3 (H3K9ac) and acetylation of lysine 27 on histone H3 (H3K27ac). Whereas H3K9ac did not decrease, we found a reproducible decrease of H3K27ac by activation of Gal4-PAX6 in this assay (**Figure 1e**). This is in accordance with the recent observations linking PAX6 to class I HDACs such as HDAC1^36^.

Importantly, we found a small but reproducible Gal4-PAX6-mediated decrease in H3K4me3 levels in this assay (**Figure 1e**). This result suggests that PAX6 may act as a transcriptional repressor at least in part by recruiting an H3K4me3 demethylase.

### KDM5C occupancy is found both on H3K4me3–positive and –negative genes

As the results from the transcriptional assays suggested that Pax6 could be acting as a repressor of transcription at least in part by bringing in H3K4me3 demethylase activity, we pursued additional *in silico* analysis of KDM5C, the most prevalent H3K4 demethylase in the developing nervous system cells^42^. This analysis revealed 7,715 KDM5C peaks (**Figure 2a**) in neurons isolated from E16.5 mouse embryonic cortices and harvested after 10 days in vitro culture ^37^, and further analysis of the most significant Reactome terms demonstrated that the most pivotal class of genes occupied by KDM5C was associated with neuronal systems (over represented p-value 3.25E-12) (**Figure 2b**).

**Figure 2:**
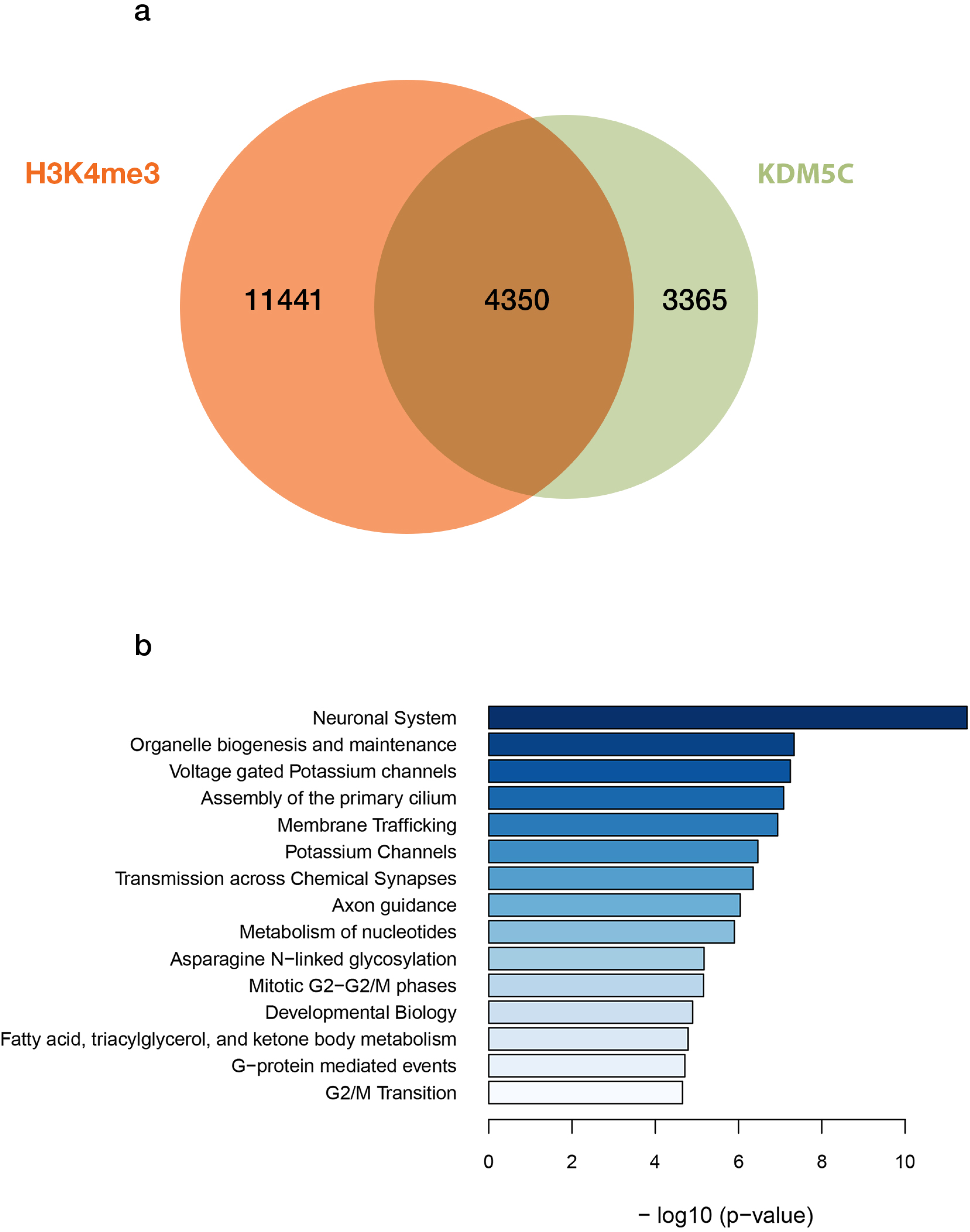
KDM5C is binding to H3K4me3-positive as well as -negative genes. (a) Analysis of ChIP-Seq experiments demonstrated KDM5C peaks both at H3K4me3-positive and -negative regions. (b) Top Reactome terms for KDM5C peaks. The most prevalent term was “neuronal system”.

Surprisingly, we found that 4,350 KDM5C peaks overlapped with the H3K4me3-positive regions detected from the embryonic forebrain ChIP-Seq (**Figure 2a**). This is interesting due to the presumed H3K4 demethylase activity of KDM5C. Nevertheless, the analysis revealed that 3,365 KDM5C peaks were associated with low H3K4me3 levels (**Figure 2a**), in accordance with the enzymatic function of KDM5C. We concluded from this analysis that KDM5C can bind to both H3K4me3-positive and -negative regions.

### Pax6 and KDM5C co-occupies 177 peaks on 131 genes mostly associated with brain development and Notch signaling

We next combined our analyses of Pax6 and KDM5C occupancies with H3K4me3 levels. This analysis revealed seven combinations of occupancies where it should be noted that the vast majority of Pax6 and KDM5C peaks did not overlap. Still 662 Pax6/KDM5C peaks were found to be present on H3K4me3-positive regions (**Figure 3a**). Most interesting in the context of the present study was however the genes that were co-occupied by Pax6 and KDM5C on regions with low H3K4me3. Here we discovered a subset of 177 peaks on 131 different genes that were co-occupied by Pax6 and KDM5C on H3K4me3-regions.

**Figure 3:**
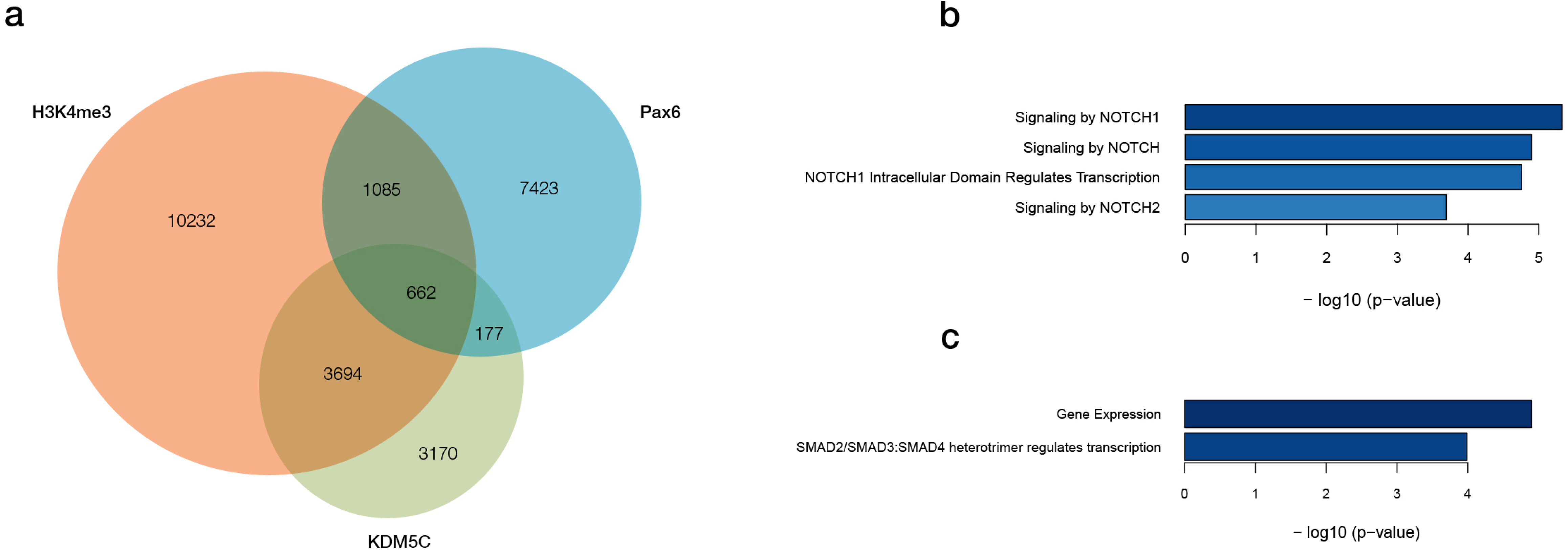
A subset of peaks where Pax6 and KDM5C overlapped with H3K4me3-negative regions was found in the proximity of genes associated with Notch signaling. (a) Analysis of Pax6, KDM5C, and H3K4me3 peaks revealed several interesting subpopulations of genes. Notably 177 peaks were found where Pax6 and KDM5C overlapped at regions with low H3K4me3 levels, but it should be noted that the vast majority of Pax6 peaks did not overlap with KDM5C or H3K4me3-positive regions. (b) The top Reactome terms for Pax6+/KDM5C+/H3K4me3-regions were found to be associated with Notch signaling.

When analyzing the biological function associated with the overlapping peaks using the Reactome terms, we made the intriguing finding that a substantial part of these 131 genes were associated with Notch signaling pathway (**Figure 3b**) (over represented p-value 4.68E-06). In contrast, when analyzing the 662 peaks of Pax6 and KDM5C co-occupancy on H3K4me3-positive regions, the top Reactome terms were *per contra* “gene expression” and BMP signaling (**Figure 3c**) (over represented p-value 1.25E-05 and 1.03E-04). These results suggest that Notch signaling factors may be specifically regulated by Pax6 and KDM5C.

### Pax6 and KDM5C peaks show distinct patterns in relation to the H3K4me3 landscape on DLL1, DLL4, and HES1 genes

We next analyzed the Pax6/KDM5C/H3K4me3-peaks in detail on specific genes involved in Notch signaling. DLL1 is a ligand of the Notch receptor, and its gene was found to contain multiple PAX6 peaks of which one overlapped with KDM5C (**Figure 4a**). The Pax6/KDM5C-overlapping peaks were flanked by high H3K4me3 levels, yet there was a small overlap of binding in a short region with low H3K4me3 levels (**Figure 4a**). DLL4 is another ligand of the Notch receptor. We could only detect one Pax6 peak in the vicinity of its gene but it overlapped with KDM5C in an H3K4-negative region of the gene, in contrast to a second KDM5C-peak that overlapped with an H3K4me3-positive site but no Pax6 binding (**Figure 4b**).

**Figure 4:**
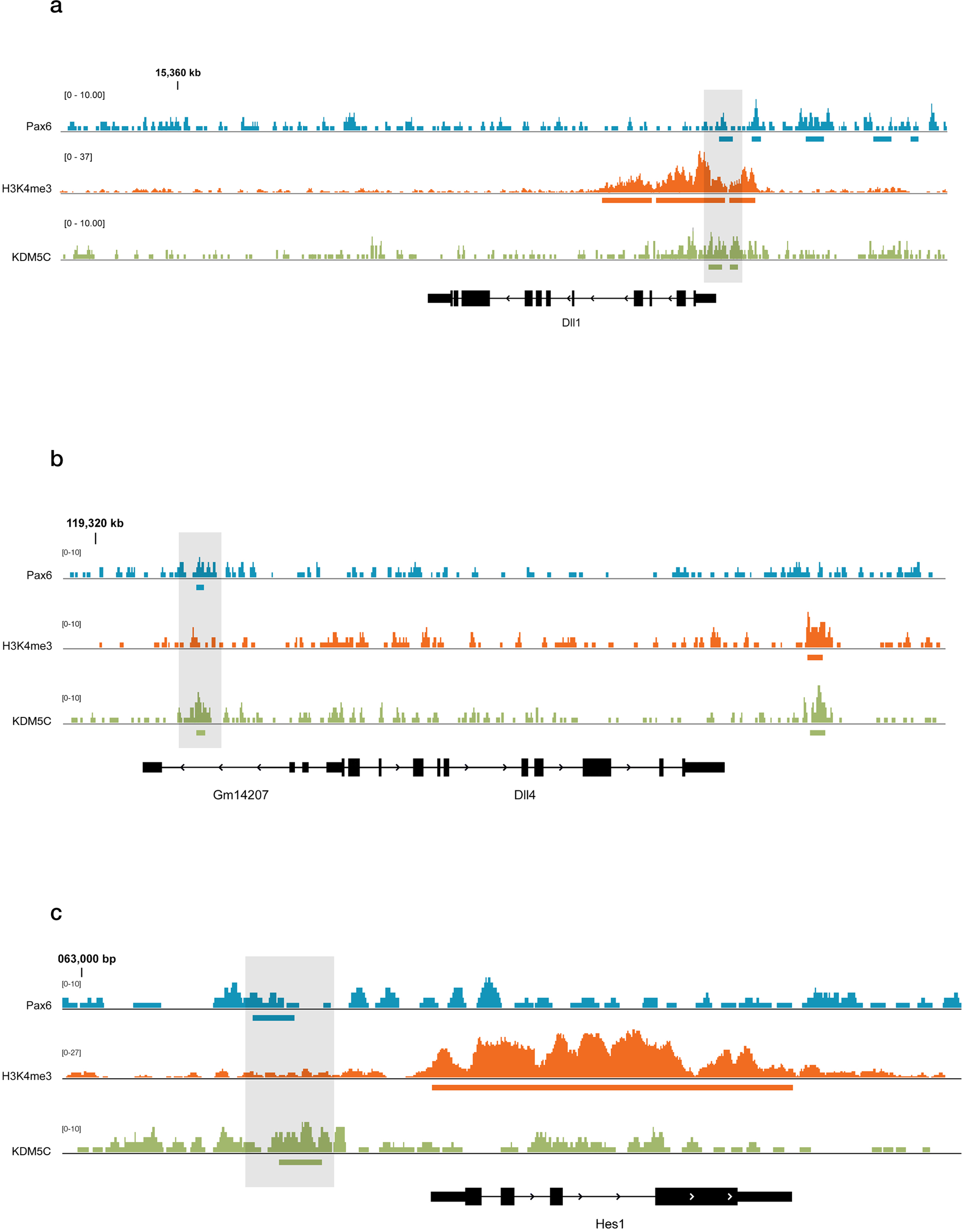
The H3K4me3 landscapes in the vicinity of the Pax6/KDM5C peaks at *Dll1, Dll4*, and *Hes1* genes display major differences. There were major differences in levels of H3K4me3 in the vicinity of the Pax6+/KDM5C+ peaks close to the genes encoding *Dll1* (a), *Dll4* (b), and *Hes1* (c). Shaded areas highlight regions with enriched Pax6 and KDM5C peaks, and low H3K4me3. Horizontal bars mark significant peaks of Pax6 (blue), KDM5C (green), and H3K4me3 (orange).

Thirdly, we analyzed the Pax6/KDM5C/H3K4me3-landscape of the gene encoding the Notch-associated transcription factor HES1. This gene showed a marked overlap of Pax6/KDM5C-overlapping peaks in an H3K4me3-negative region, whereas downstream of this site, we found very high levels of H3K4me3 (**Figure 4c**). In summary, whereas we can confirm the overlap of Pax6 and KDM5C peaks on Notch signaling-related genes, the peaks showed variability in character and the gene regions displayed major variations in the H3K4me3 landscape, hinting at possible differences in gene regulatory mechanisms.

### RNA knockdown of PAX6 and KDM5C in human iPS-derived neural progenitor cells result in differential gene expression levels of DLL1 and DLL4 but not HES1

Lastly, to investigate whether the detected Pax6/KDM5C peaks in neural progenitors on Notch signaling-associated genes were of relevance for regulation of gene expression, we developed specific siRNAs to perform combined RNA knockdown of PAX6 and KDM5C and compared this to control (scrambled) siRNA in human iPS-derived neural progenitors (hNPs) to analyze the effects of PAX6 /KDM5C knockdown (PAX6 siRNA (*p*=0.0005) and KDM5C siRNA (*p*=0.0002)) on the expression levels of *DLL1, DLL4*, and *HES1*.

We did not detect any significant differences in the expression of *HES1* after PAX6/KDM5C RNA knockdown in hNPs compared to control (**Figure 5**). As noted (**Figure 4c**), the *HES1* gene region was very rich in H3K4me3, suggesting that PAX6/KDM5C may not be dominant repressive regulators of the expression of this gene. When analyzing the expression of *DLL1*, we found that the gene expression decreased (p=0,0040) after RNA knockdown of PAX6/KDM5C in hNPs compared to control.

**Figure 5:**
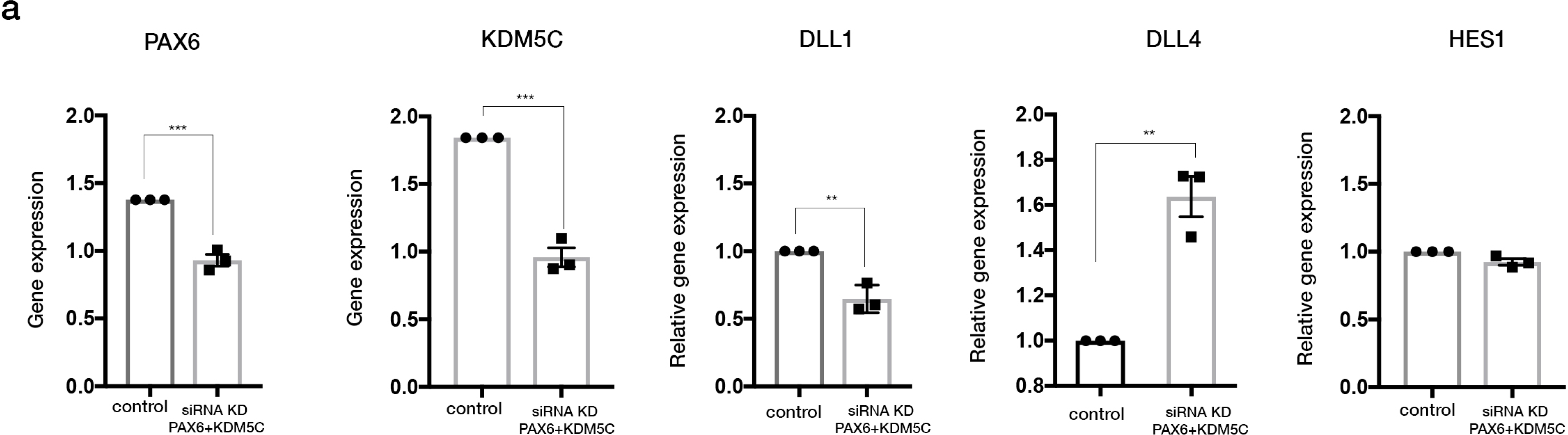
Effects on gene expression levels of *DLL1, DLL4*, and *HES1* after RNA knockdown of PAX6 and KDM5C in human iPS-derived neural progenitors. (a) The RNA knockdown of PAX6 and KDM5C (left) after specific siRNA administration was significant (PAX6 siRNA (n=3, \*\*\**p*=0.0005; unpaired, two-tailed t-test) and KDM5C siRNA (n=3, \*\*\**p*=0.0002; unpaired, two-tailed t-test). There was a significant decrease in *DLL1* (middle) gene expression (n=3, \*\**p*=0.0040; unpaired, two-tailed t-test), a significant increase in *DLL4* gene expression (n=3, \*\**p*=0.0020; unpaired, two-tailed t-test), and no significant difference in HES1 mRNA levels (right) after the simultaneous knockdown of PAX6 and KDM5C mRNAs in hNPs. Error bars represent ± SEM.

Lastly, we analyzed the gene expression levels of *DLL4* after RNA knockdown of PAX6/KDM5C in the hNPs, and found a robust increase (p=0,0020) in the mRNA levels compared to control (**Figure 5**).

In summary, our results demonstrate that there is a relatively small subset of developmentally critical genes, including Notch signaling factors, co-occupied by PAX6 and KDM5C on H3K4me3-negative regions, and that these two factors exert essential control of proper expression of some of these genes.

## DISCUSSION

Here we have revealed that PAX6 and KDM5C co-occupy and regulate a subset of developmentally critical genes associated with Notch signaling. Considering the fundamental role of Notch signaling in neural development, we suggest that this regulatory mechanism may play a role in pathologies associated with aberrant PAX6 and/or KDM5C activity.

This is to our knowledge the first link between KDM5C and regulation of Notch signaling factors, and it will be of immediate interest to investigate whether effects on Notch signaling could be involved in the consequences of aberrant KDM5C function^37,43^. In contrast, Pax6 has been linked to Notch signaling in various contexts, but that Pax6 is binding to regions close to almost all known factors directly involved in Notch signaling has not been previously understood. In fact, the analysis revealed PAX6 binding close to all ligands and all receptors, except NOTCH3, as well as more distant members, such as DLK1 (G.G and O.H., unpublished observations). It is likely that this has not been shown before due to the large number of genes to which PAX6 is binding and directly regulating^31^, and it is not until analyzing the co-occupied regions with KDM5C that this subpopulation of genes is revealed.

PAX6 has previously been linked to chromatin modifying and remodeling factors, such as CBP/p300 and members of the SWI/SNF and BAF complexes (cf. ^44^, reviewed in ^31^). Less is however known about the mechanisms underlying transcriptional repression by Pax6. Our findings show that Pax6 and KDM5C co-occupy regions with low H3K4me3 but many peaks of PAX6 found at regions with low H3K4me3 were not occupied by KDM5C and thus presumably not active (**Figure 3a**). Therefore, alternative mechanisms of repression should be considered. It was recently shown that Pax6 binds to the histone deacetylase HDAC1 with functional implications for lens development^36^. HDAC1 is however not ubiquitously expressed in neural progenitors ^11^, and in the hNPs used in the present study, another class I HDAC, HDAC2, is the most prominently expressed (M. Lam and A. Falk, personal communications). Class I HDACs such as HDAC1 and HDAC2 can be members of the so called NURD complex that among other things regulate H3K27 acetylation, and indeed we found in the transcriptional assays that PAX6 overexpression resulted in a decrease in H3K27 acetylation (**Figure 1**). Future studies will aim at elucidating whether PAX6 can interact with other class I HDACs and possibly the NURD complex in hNPs to regulate the H3K27 acetylation.

As demonstrated in Figure 5, simultaneous RNA knockdown of PAX6 and KDM5C did not result in an increased gene expression of all genes assessed. At a first glance, this may seem like a contradiction, but the variations in the chromatin landscape surrounding the PAX6/KDM5C binding sites must be taken into account. Thus, in the case of *DLL1*, there are multiple PAX6 binding sites in the gene but only one overlapping peak with KDM5C (**Figure 4a**). It may therefore be hypothesized that the decrease in *DLL1* gene expression was a net result by knocking down PAX6 not only as a repressor but also as an activator, and that PAX6 may be predominantly an activator of the *DLL1* gene. Further, the chromatin landscape in the vicinity of the PAX6/KDM5C binding site at the *HES1* gene was very rich in H3K4me3 (**Figure 4c**), pointing to an ancillary role for PAX6/KDM5C in the regulation of *HES1* gene expression.

When analyzing the transcriptional outcome, it should be noted that the chromatin is organized in a 3-dimensional manner, and it is the nanoorganization - or nanoarchitecture - in coordination with the chemical milieu that will provide the optimal conditions for transcriptional regulation. Such parameters are extremely dynamic, especially in progenitor cells. A common model of the outcome of chromatin modifications is the “gas and brake” of a car, where H3K4me3 is compared to the gas and H3K27me3 to the brake. The model is of course knowingly a simplification, but staying with the car analogy, a simplified model for transcriptional regulation by chromatin modifications could rather be compared to a “clutch”, as previously proposed^11^, where the state of transcription and nanoarchitecture will influence the effect of changes in transcription and chromatin modifying factor levels. As the surrounding chemical architecture will determine the overall transcriptional activity, the effect(s) of simple RNA knockdown or overexpression will depend on the overall enhancer/promoter activity, and the effects will be different depending on the baseline of the transcription of the gene assessed, the number of transcription factor binding sites, the nature of the modification etc. To test this hypothesis, future studies should be utilizing controlled genetic and epigenetic editing in neural progenitors in a 3D system. Our findings reveal a relatively small population of genes in neural progenitors with low levels of H3K4me3 that are co-occupied by Pax6 and KDM5C, and a subset of those are genes encoding factors involved in Notch signaling.

## METHODS

### Bioinformatic analysis

Relevant and publicly available datasets were identified via literature search and sratools/2.8.0 was used to download raw sequencing reads in fastq format from The Sequence Read Archive^37,38^. Best practices ChIP-seq data processing and analyses as outlined by ENCODE consortium were followed to analyse the data. Briefly, TruSeq adapters were removed from the data as well as poor quality reads using trimmomatic/0.32. Alignment against reference GRCm38.p5 genome was done with bowtie/1.1.2 suppressing all multiple alignments. Duplicated reads and/or reads overlapping with the blacklisted genomics regions with artificially high signals were removed with picard/2.0.1 (https://broadinstitute.github.io/picard/.). Peak-independent quality metrics, including strand cross-correlation, were calculated with phantompeakqualtools/1.1 and deepTools/1.1.2 to assess quality of ChIP-seq libraries (strength of enrichment). Peaks were called with MACS/2.1.0^45^ using extension size as calculated from strand cross-correlation (200 and 190 for S1 and S3 respectively). Peaks called were pooled across replicates where feasible using BEDTools/2.26.0. Annotation to the closest genes and overlapping analyses of ChIP-seq peaks were done using R/Bioconductor package ChIPpeakAnno^46^. Genes associated with peaks were analysed for over-represented GO and reactome terms using R/ Bioconductor package goseq^47^.

### GAL4 plasmids and DNA cloning

GAL4–*PAX6* was cloned into the pcDNA5/FRT/TO expression vector by standard procedures and subsequently used for recombination into the genomic DNA of TREx 293 Flp-In cells (Invitrogen) as described in details previously^41^. The GAL4-*PAX6* constructs were sequence-verified by DNA sequencing. GAL4–EZH2 (control) was a kind gift from Dr. Kristian Helin.

### Cell culture neuroepithelial stem cells

The generation of AF22, a control iPS cell line, has been described elsewhere^48^. Briefly, to culture long-term self-renewing neuroepithelial-like stem cells or neural progenitors (hNPs), hiPS cell colonies were induced to differentiate to neural tissue and hNPs cultures were captured and maintained by passaging them at the ratio of 1:3 every third day essentially as previously described^48^. Briefly, hNPs cultures were maintained in hNPs medium (DMEM/F-12, GlutaMAX containing N_2_ supplement, FGF2 (10 ng ml^-1^), B27 (0.1%), Pen/Strep (all from Invitrogen), and EGF (10 ng ml^-1^; Peprotech, Rocky Hill, NJ, USA)), and harvested using TrypLE-Express (Gibco) for passage into 0.1 mg poly-L-ornithine (Sigma-Aldrich) and 1 μg ml^-1^laminin L2020 (Sigma-Aldrich) coated (PLO/L-coated) plates at 1:3 ratio.

### Chromatin Immunoprecipitation (ChIP-qPCR)

Chromatin immunoprecipitation (ChIP) was performed using HighCell# ChIP kit (Diagenode) and performed according to the manufacturer’s protocol. 6,3 micrograms of the specific antibody (H3K4me3, H3K9ac and H3K27ac all Rabbit mAb from Cell Signaling Technology) were used in each IP. Purified, eluted DNA and 1% DNA Input was analyzed by qPCR using Platinum™ SYBR™ Green qPCR SuperMix-UDG (ThermoFisher Scientific) together with site-specific primers. The analyzes were performed in triplicates per each biological sample using the standard curve method to account for potential differences in primer efficiency, therefore relative occupancy for a specific antibody in different regulatory regions could be compared. Statistical analyzes were performed using GraphPad Prism (version 7.0, GraphPad Software, La Jolla, CA, USA).

### RNA knockdown

For siRNA-mediated knockdown of PAX6 RNA, KDM5C RNA, and control, Amaxa Nucleofector™ Kits for Human Stem Cells (Lonza) was used according to the manufacturer’s protocol. ON-TARGET plus SMARTpool siRNA for PAX6, KDM5C, and control were obtained from Dharmacon. Following nucleofection, hNPs were seeded in 6 well plates in hNPs media under proliferative conditions, as described in the cell culture method section. The results were analyzed 48h later by RT-qPCR.

### RT-qPCR

Total RNA was extracted using RNeasy Mini Kit (Qiagen), cDNA was synthesized using High Capacity cDNA Reverse Transcription Kit (Applied Biosystems, now ThermoFisher Scientific), and RT-qPCR was performed using Platinum™ SYBR™ Green qPCR SuperMix-UDG (Invitrogen, now ThermoFisher Scientific) together with site specific primers. mRNA expression levels were normalized to the housekeeping human TATA-binding protein (TBP) mRNA (RT^2^ qPCR Primer Assays, Qiagen) in the GAL4 assay and to the human housekeeping hypoxanthine guanine phosphoribosyltransferase (HPRT) (RT^2^ qPCR Primer Assays, Qiagen) mRNA when performing siRNA analyses.

## ACKNOWLEDGEMENTS

We would like to thank members of the Hermanson lab for valuable input on the study. The authors are especially grateful to Ricardo Paap, Matti Lam, Anna Vidal, and Hannah Bruce for experimental support, and Anna Falk for the hNPs. The computations were performed on resources provided by SNIC through Uppsala Multidisciplinary Center for Advanced Computational Science (UPPMAX). The authors would like to acknowledge support from Science for Life Laboratory, the National Genomics Infrastructure, NGI, and Uppmax for providing assistance in massive parallel sequencing and computational infrastructure. The study was supported by grants from the Swedish Research Council (VR-MH), the Ming Wai Lau Centre for Reparative Medicine (MWLC), the Swedish Cancer Society (CF), and the Swedish Childhood Cancer Foundation (BCF).

## SUPPLEMENTARY FILES

**Supplementary File 1:** Detailed methods and results of the bioinformatic analysis.

**Supplementary Figure 1:** (a) Three different PAX6-GAL4 clones exerted repression in the assay in a similar range as the Polycomb-associated repressor EZH2. (b) The protein product of the GAL4-PAX6 construct was specifically induced by DOX treatment.

