## Supplementary figures and images for "Pax6 and KDM5C co-occupy a subset of developmentally critical genes including Notch signaling regulators in neural progenitors"

### Supplementary Figure 1

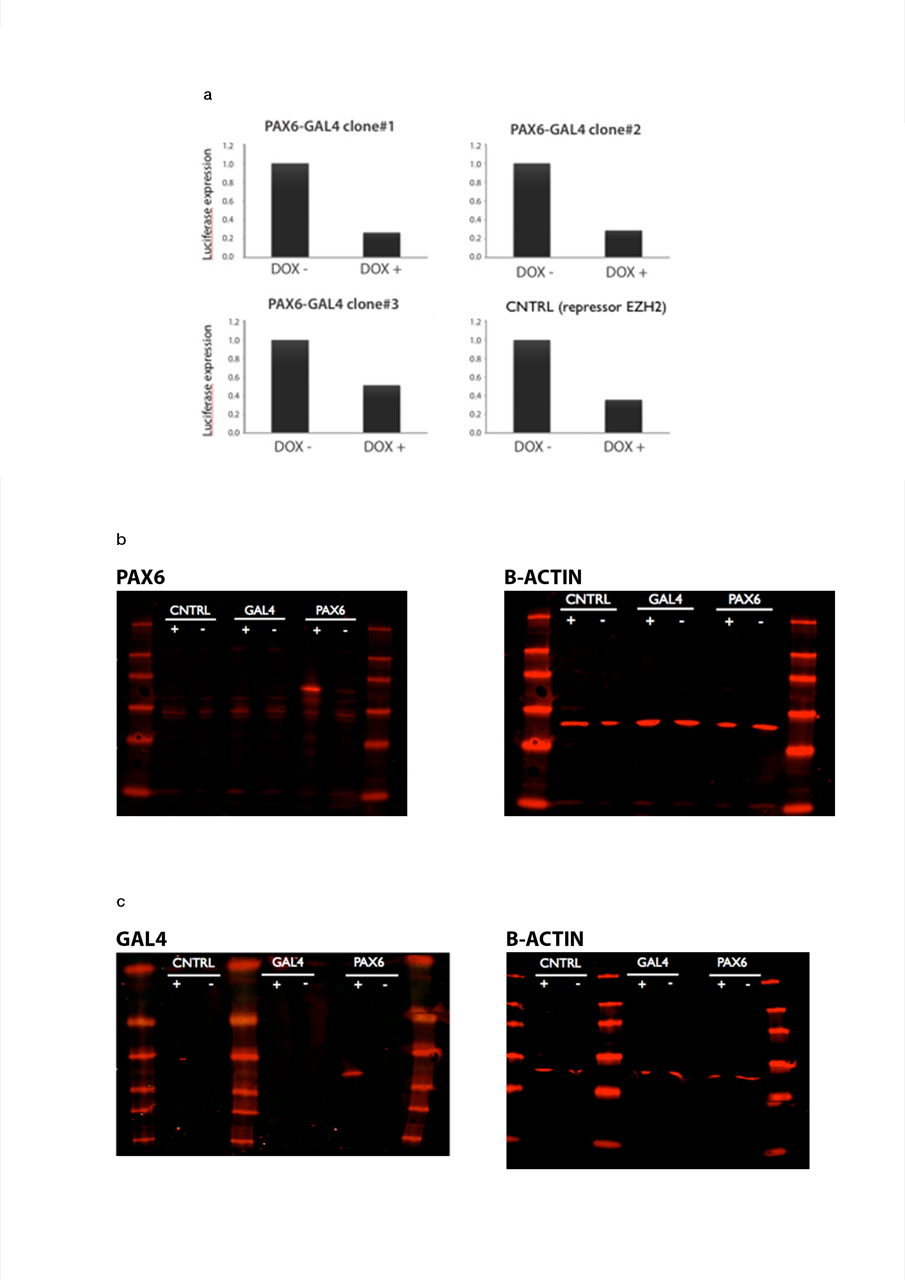
