## Supplementary File 1 for "Pax6 and KDM5C co-occupy a subset of developmentally critical genes including Notch signaling regulators in neural progenitors"

### 1 Materials and Methods

#### 1.1 Available data

**S1 & S3** A set of publicly available ChIP-seq data were identified, downloaded from GEO/SRA and analyzed, as shown in Table 1

| ID | PMID | GSE/SRR | description |
|---|---|---|---|
| S1 | PMID:26804915 | GSE61036 | KDM5c_Spikein_Input_Pooled_CN_rep2 |
| S1 | PMID:26804915 | GSE61036 | KDM5c_Spikein_SMCX_333_ChIP_KO_CN_rep2 |
| S1 | PMID:26804915 | GSE61036 | KDM5c_Spikein_SMCX_333_ChIP_WT_CN_rep2 |
| S1 | PMID:26804915 | GSE61036 | KDM5c_Spikein_Input_Pooled_CN_rep1 |
| S1 | PMID:26804915 | GSE61036 | KDM5c_Spikein_SMCX_333_ChIP_KO_CN_rep1 |
| S1 | PMID:26804915 | GSE61036 | KDM5c_Spikein_SMCX_333_ChIP_WT_CN_rep1 |
| S3 | PMID:26138486 | SRP056219 | forebrain_Pax6_ChIPseq_rep1 |
| S3 | PMID:26138486 | SRP056219 | forebrain_Pax6_ChIPseq_rep2 |
| S3 | PMID:26138486 | SRP056219 | forebrain_input_3 |
| S3 | PMID:26138486 | SRP056219 | forebrain_input_4 |
| S3 | PMID:26138486 | SRP056219 | forebrain_H3K4me3_ChIPseq |
| S3 | PMID:26138486 | SRP056219 | forebrain_input_2 |
| S3 | PMID:26138486 | SRP056219 | forebrain_input_1 |

Table 1: Selected, downloaded and analyzed GEO/SRA data

#### 1.2 Methods

- `sratools/2.8.0` was used to download raw sequencing reads, fastq files, from The Sequence Read Archive (SRA)
- `FastQC/0.11.5` [Andrews 2010] was used to check the quality on raw sequencing reads, separate per each .fastq.gz file
- reads sequenced over the two lanes were concatenated with `cat` command
- `trimmomatic/0.32` [Breese and Liu 2013] was used to remove TruSeq adapters as well as to trim and filter reads. Reads shorter than 36bp and/or with average Phred quality score lower than 18, within 4-base wide sliding window, were removed.
- alignment against reference `GRCm38.p5` genome was done with `bowtie/1.1.2` [Langmead 2011] suppressing all multiple alignments [bowtie -n 2 –best -m 1 -q -S -p 3 -e 80 -t –chunkmbs 512]
- `samtools/1.3` [Li et al. 2009] was used to convert the aligned reads in .SAM format to .BAM format as well as to sort and index the .BAM files
- `phantompeakqualtools/1.1` was used to calculate the standard cross-correlation metrics
- the blacklisted genomic regions with artificially high signals were downloaded for mm9 and lifted over to mm10 using `LiverOver` USCS Genome Browser Tools facility.
- `picard/2.0.1` [BroadInstitute] was run to remove the duplicated reads, `NGSUtils/0.5.9` [Breese and Liu 2013]to remove the regions overlapping with the blacklisted genomic regions with artificially high signals, as downloaded from `https://www.encodeproject.org/annotations/ENCSR636HFF/`
- `phantompeakqualtools/1.1` [Phantompeakqualtools] was used again to calculate the standard cross-correlation metrics and plots after removal of the duplicated reads and blacklisted genomic regions
- `deepTools/1.1.2` [Ramírez et al. 2014] was run to obtain coverage tracks normalized to 1xcoverage, with using 2.15e9 as an estimated effective genome size, to obtain ChIP-seq cumulative enrichment (fingerprint) and to assess overall similarity between libraries, spearman and pearson based clustering heatmaps in bin mode (with bin size of 5000 bp)

- `MACS/2.1.0` [Feng et al. 2012] was used to call peaks, with 200 (S1) and 190 (S3) as computed fragment length from the cross-correlation analysis [-q 0.01 –nomodel –extsize 200/190]
- `BEDOPS/2.4.3` [Neph et al. 2012] was used to prepare a combined list of peaks and similarity between libraries in BED mode was re-calculated using `deepTools/1.1.2`
- peaks called were pooled across replicates where feasible using `BEDTools/2.26.0`
- annotation to the closest genes and overlapping analyses of ChIP-seq peaks were done using R / Bioconductor package `ChIPpeakAnno` [Zhu et al. 2010] item genes associated with peaks were analysed for over-represented GO and reactome terms using R / Bioconductor package `goseq` [Young et al. 2010]. R session info listing all packages and dependencies can be found at the end of this document.

### 2 Results: S1

#### 2.1 Peak-independent quality metrics

##### 2.1.1 Cross-correlation

| | fragLength | NSC | RSC | QS |
|---|---|---|---|---|
| SRR2984333 | 195,320,340 | 1.06 | 2.25 | 2 |
| SRR2984334 | 195,320,340 | 1.06 | 2.25 | 2 |
| SRR2984335 | 195,320,340 | 1.06 | 2.25 | 2 |
| SRR2984336 | 195,320,340 | 1.06 | 2.25 | 2 |
| SRR2984337 | 195,320,340 | 1.06 | 2.25 | 2 |
| SRR2984338 | 195,320,340 | 1.06 | 2.25 | 2 |

Table 2: Cross-correlation measures

| | fragLength | NSC | RSC | QS |
|---|---|---|---|---|
| SRR2984333 | 195,205,355 | 1.05 | 1.94 | 2 |
| SRR2984334 | 195,205,355 | 1.05 | 1.94 | 2 |
| SRR2984335 | 195,205,355 | 1.05 | 1.94 | 2 |
| SRR2984336 | 195,205,355 | 1.05 | 1.94 | 2 |
| SRR2984337 | 195,205,355 | 1.05 | 1.94 | 2 |
| SRR2984338 | 195,205,355 | 1.05 | 1.94 | 2 |

Table 3: Cross-correlation measures after removing duplicated reads and blacklisted genomic regions

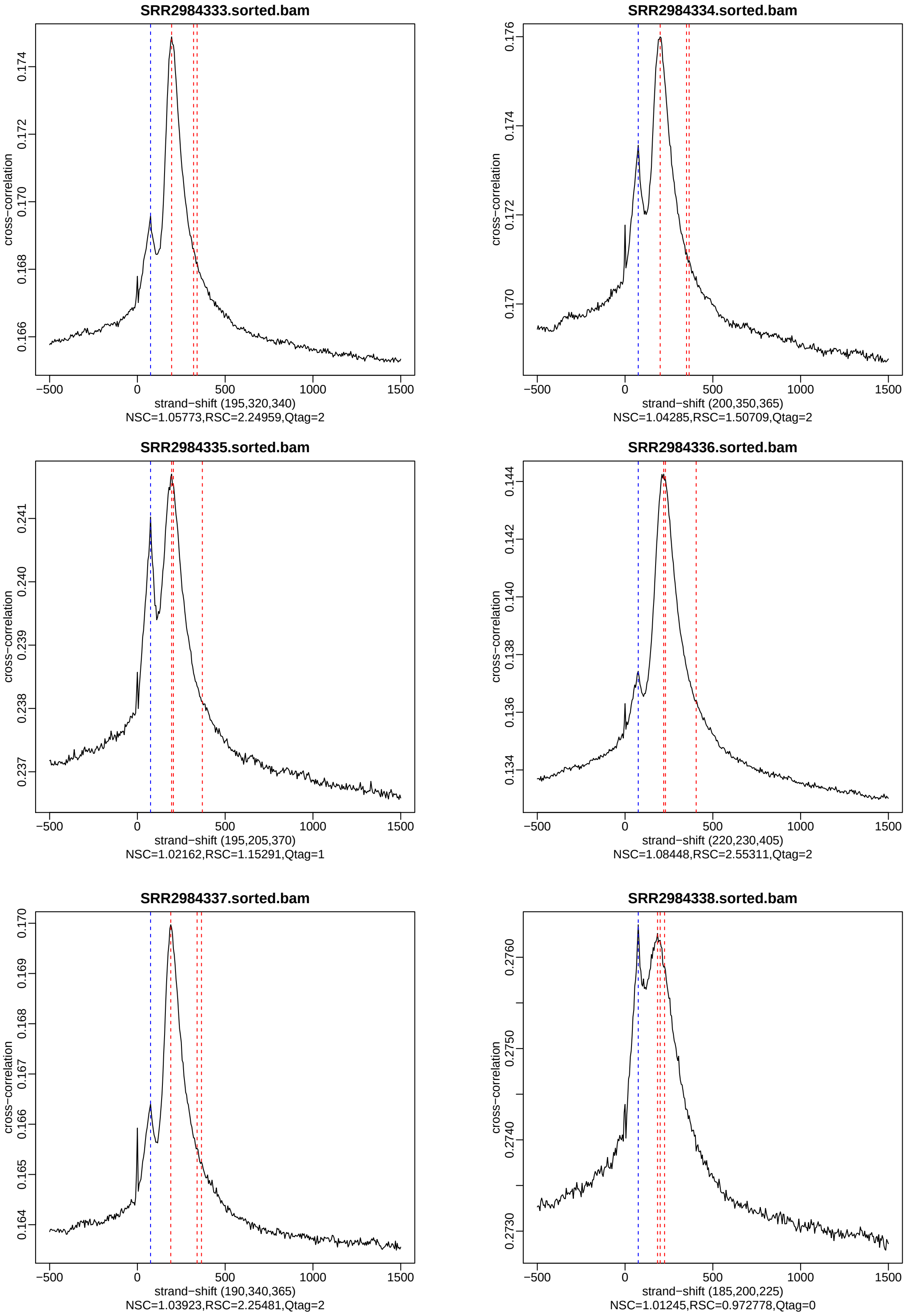


Figure 1: Cross-correlation profile for ChIP (SRR2984333, SRR2984334, SRR298436, SRR2984337) and input (SRR2984335, SRR2984338)

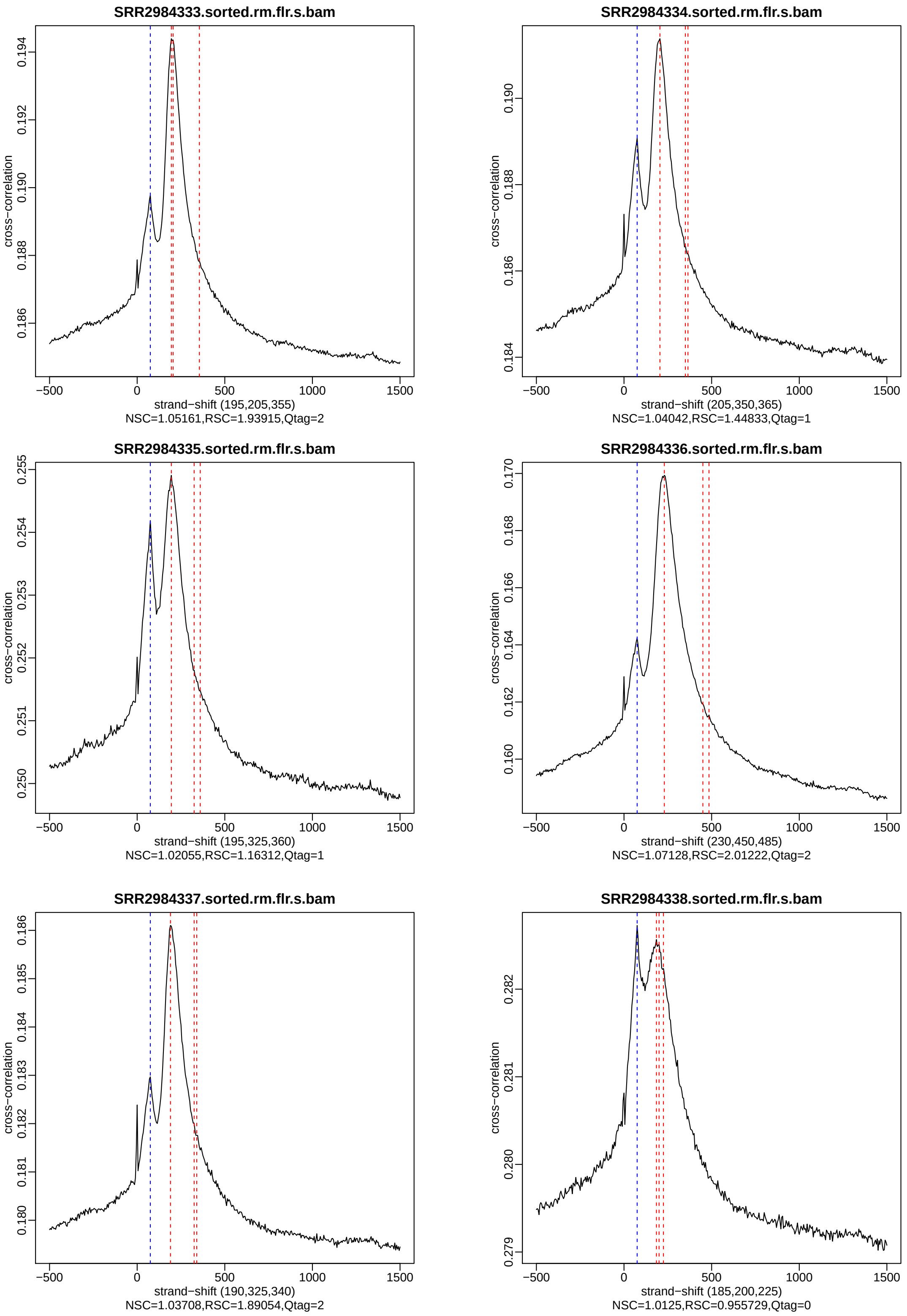


Figure 2: Cross-correlation profile for ChIP (SRR2984333, SRR2984334, SRR298436, SRR2984337) and input (SRR2984335, SRR2984338) after removing duplicated reads and blacklisted genomic regions

##### 2.1.2 Cumulative enrichment

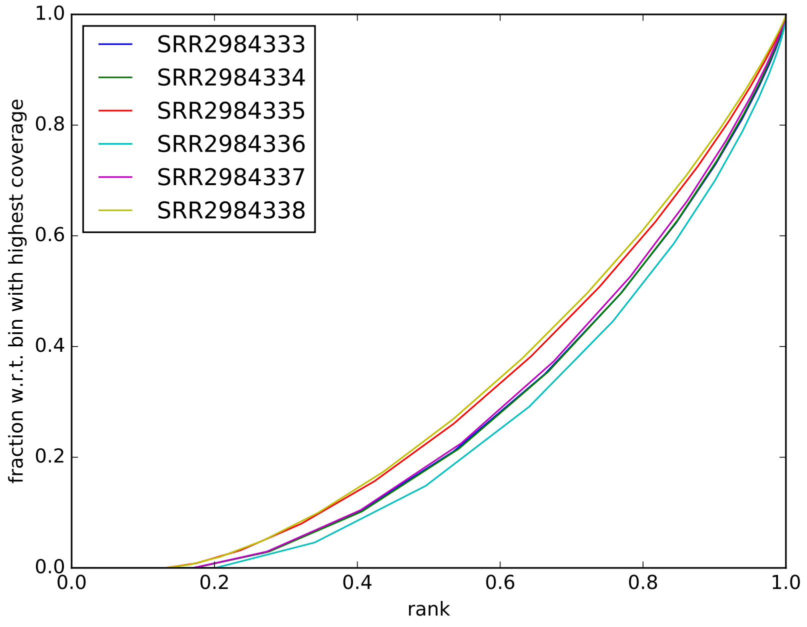


Figure 3: Cumulative enrichment in all libraries

##### 2.1.3 Library clustering by similarity (BIN mode)

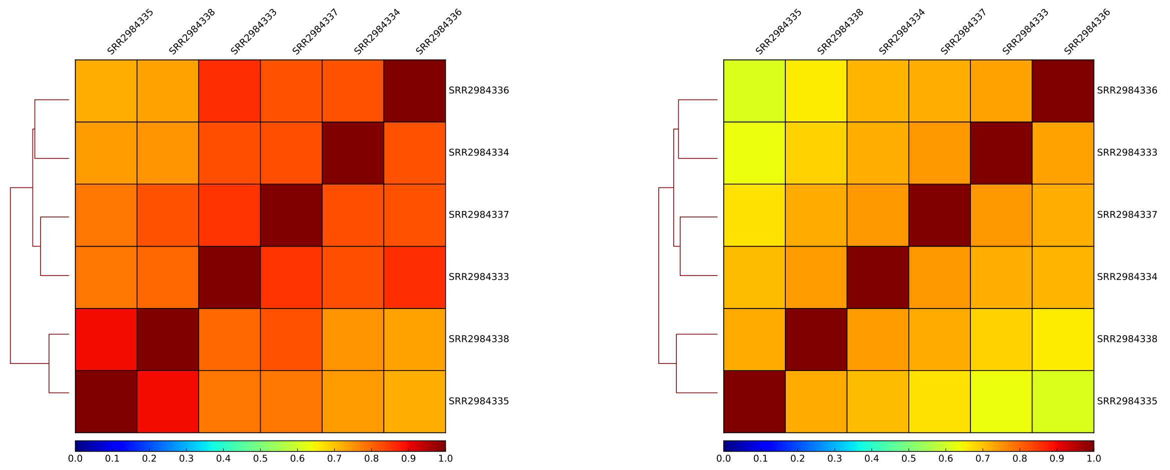


Figure 4: Pearson (left) and Spearman (right) correlation of signal in all libraries

#### 2.2 Peak calling

##### 2.2.1 Peaks statistics

| library | peaks |
|---|---|
| SRR2984333_vs_SRR2984335 | 1332 |
| SRR2984333_vs_SRR2984338 | 2864 |
| SRR2984334_vs_SRR2984335 | 3301 |
| SRR2984334_vs_SRR2984338 | 3851 |
| SRR2984336_vs_SRR2984335 | 9444 |
| SRR2984336_vs_SRR2984338 | 15595 |
| SRR2984337_vs_SRR2984335 | 15639 |
| SRR2984337_vs_SRR2984338 | 15672 |

Table 4: Number of peaks detected

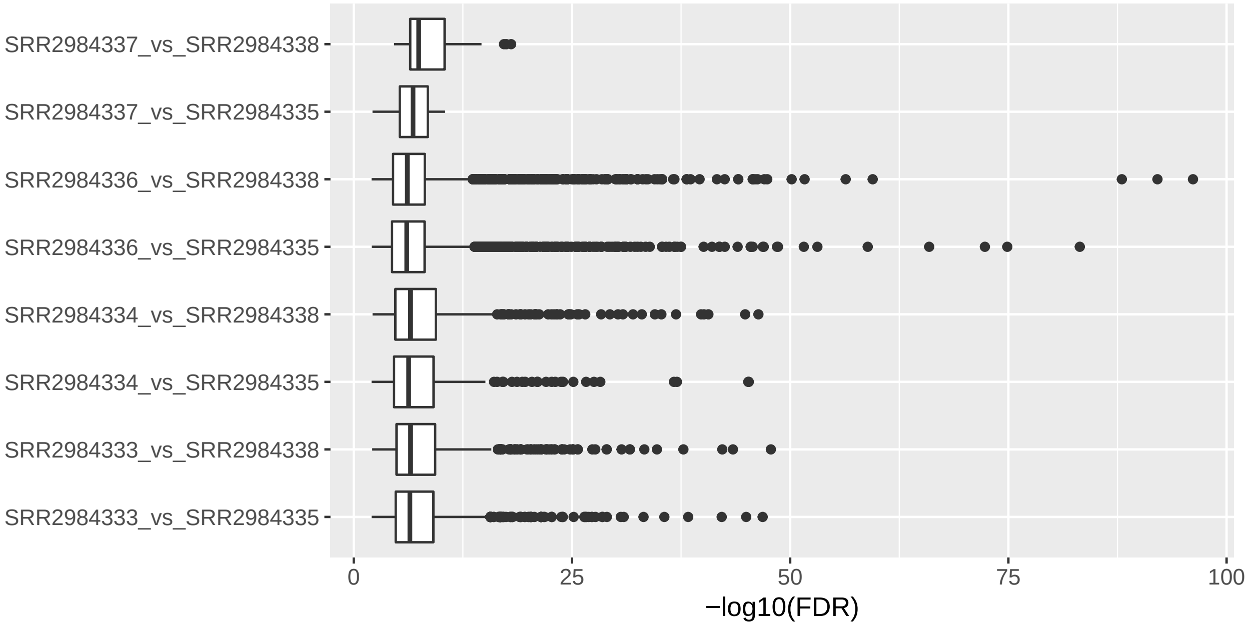


Figure 5: Boxplots of q values for the peaks detected

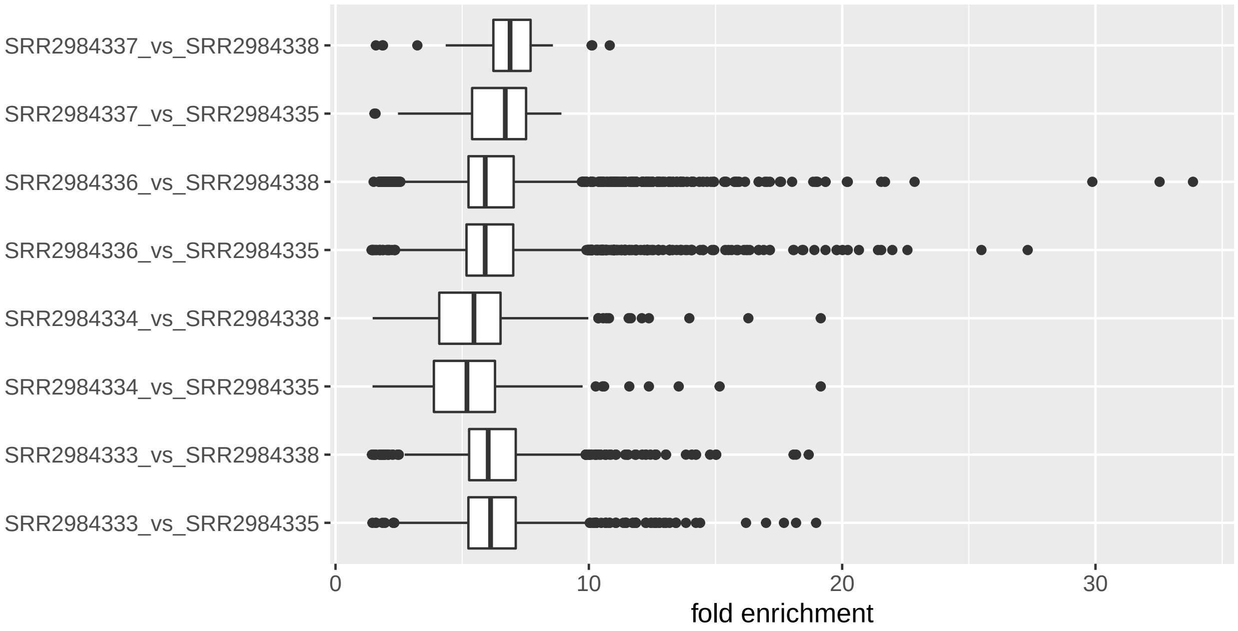


Figure 6: Boxplots of fold change values of peaks detected

###### 2.2.2 Library clustering by similarity (BED mode)

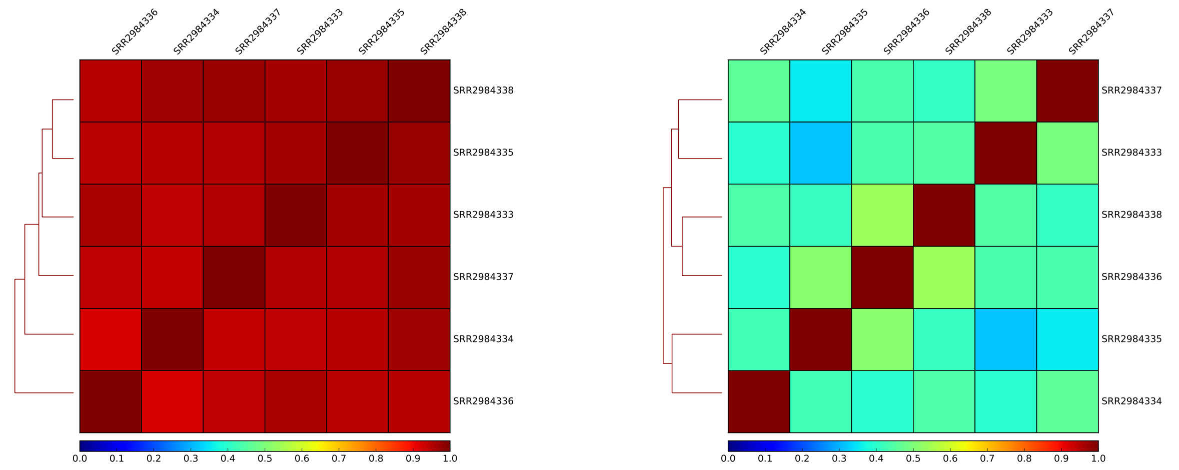


Figure 7: Pearson (left) and Spearman (right) correlation of signal in all libraries, in BED mode, based on the identified peaks regions.

### 3 Results: S3

#### 3.1 Peak-independent quality metrics

###### 3.1.1 Cross-correlation

| | fragLength | NSC | RSC | QS |
|---|---|---|---|---|
| SRR1916963 | 185,200,210 | 1.02 | 1.00 | 0 |
| SRR1916964 | 185,200,210 | 1.02 | 1.00 | 0 |
| SRR1916965 | 185,200,210 | 1.02 | 1.00 | 0 |
| SRR1916966 | 185,200,210 | 1.02 | 1.00 | 0 |
| SRR1916971 | 185,200,210 | 1.02 | 1.00 | 0 |
| SRR1916977 | 185,200,210 | 1.02 | 1.00 | 0 |
| SRR1916979 | 185,200,210 | 1.02 | 1.00 | 0 |

Table 5: Cross-correlation measures

| | fragLength | NSC | RSC | QS |
|---|---|---|---|---|
| SRR1916963 | 185,200,215 | 1.02 | 0.94 | 0 |
| SRR1916964 | 185,200,215 | 1.02 | 0.94 | 0 |
| SRR1916965 | 185,200,215 | 1.02 | 0.94 | 0 |
| SRR1916966 | 185,200,215 | 1.02 | 0.94 | 0 |
| SRR1916971 | 185,200,215 | 1.02 | 0.94 | 0 |
| SRR1916977 | 185,200,215 | 1.02 | 0.94 | 0 |
| SRR1916979 | 185,200,215 | 1.02 | 0.94 | 0 |

Table 6: Cross-correlation measures after removing duplicated reads and blacklisted genomic regions

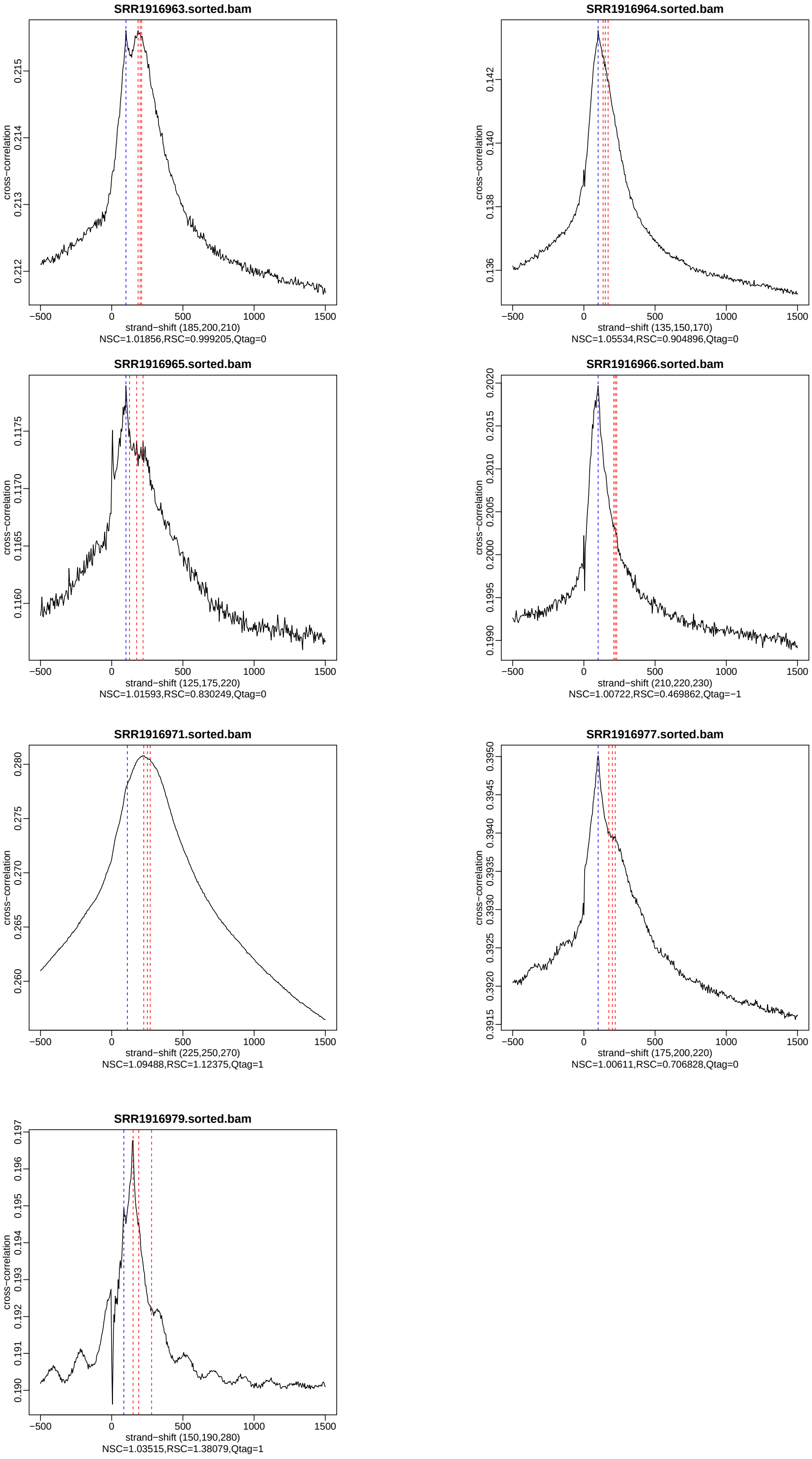


Figure 8: Cross-correlation profile for ChIP (SRR1916963, SRR1916964, SRR1916971) and input (SRR1916965, SRR1916966, SRR1916977, SRR1916979)

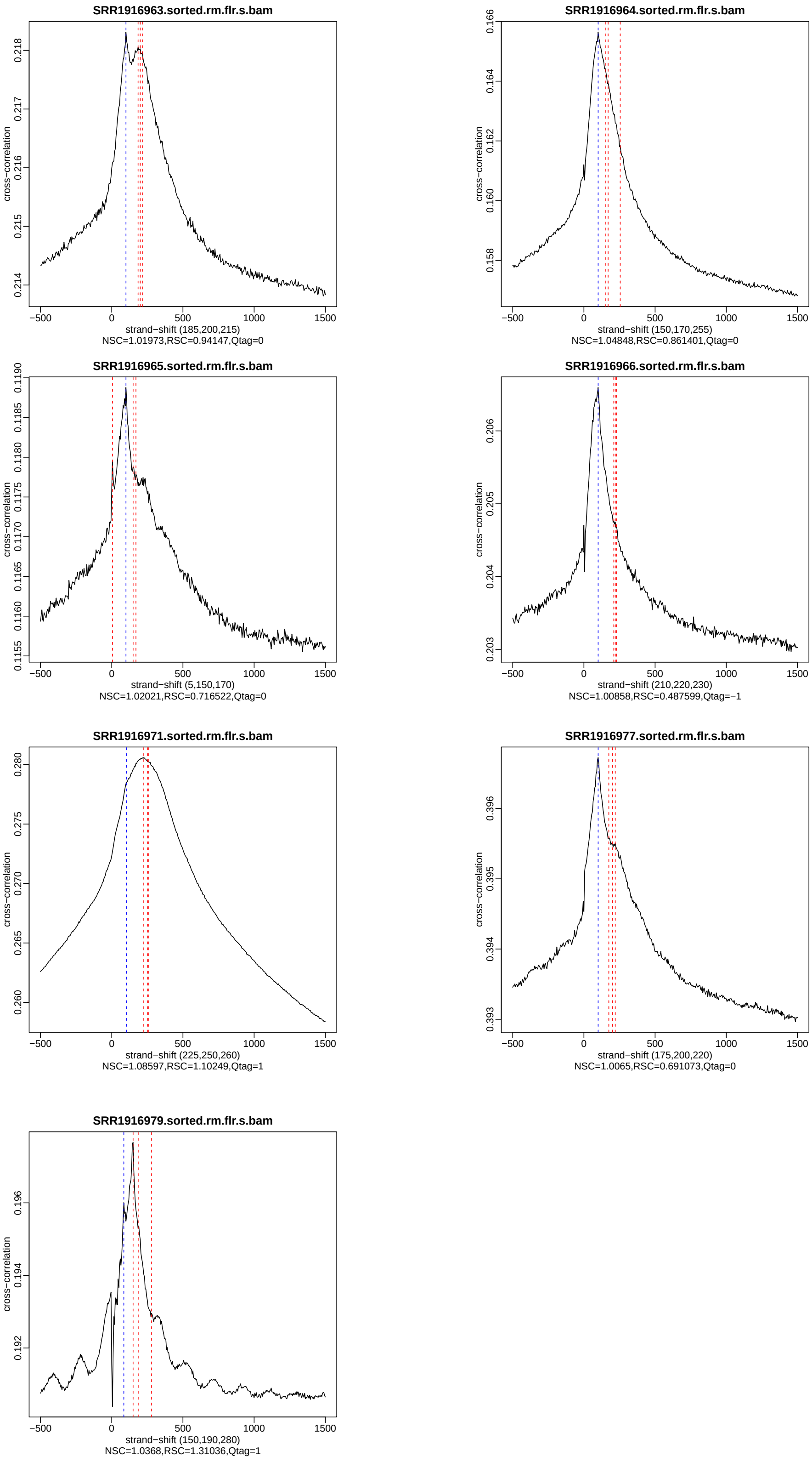


Figure 9: Cross-correlation profile for ChIP (SRR1916963, SRR1916964, SRR1916971) and input (SRR1916965, SRR1916966, SRR1916977, SRR1916979) after removing duplicated reads and blacklisted genomic regions

##### 3.1.2 Cumulative enrichment

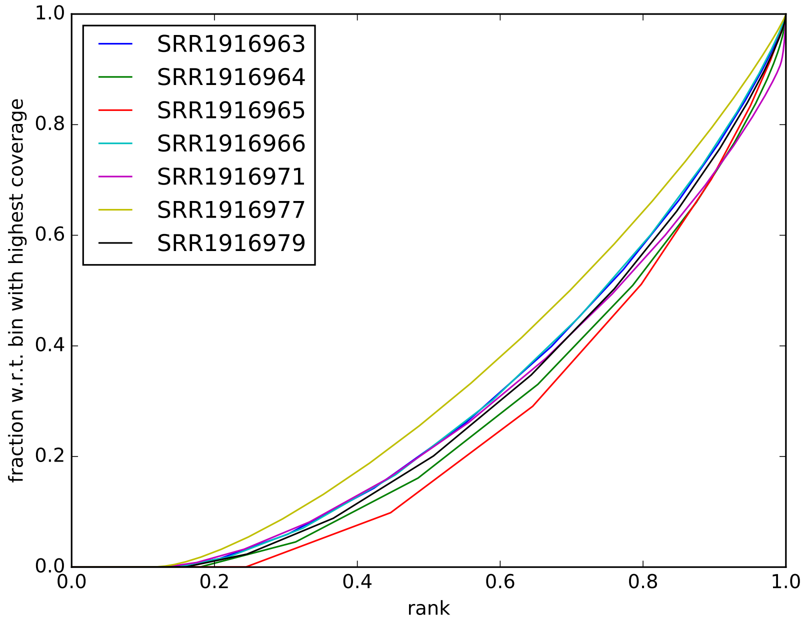


Figure 10: Cumulative enrichment in all libraries

##### 3.1.3 Library clustering by similarity (BIN mode)

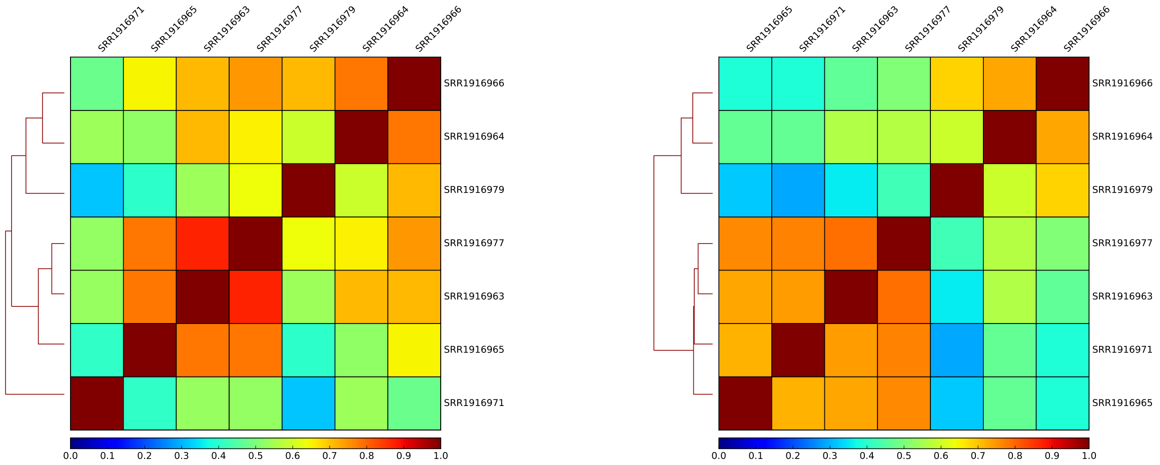

Figure 11: Pearson (left) and Spearman (right) correlation of signal in all libraries

#### 3.2 Peak calling

Peak-independent quality metrics, Figures 8, 9, 10 and 11, indicate that the best input library replicate is SRR1916977 and therefore it was used for peak calling.

##### 3.2.1 Peaks statistics

| library | peaks |
|---|---|
| SRR1916963_vs_SRR1916977 | 2247 |
| SRR1916964_vs_SRR1916977 | 11368 |
| SRR1916971_vs_SRR1916977 | 27242 |

Table 7: Number of peaks detected

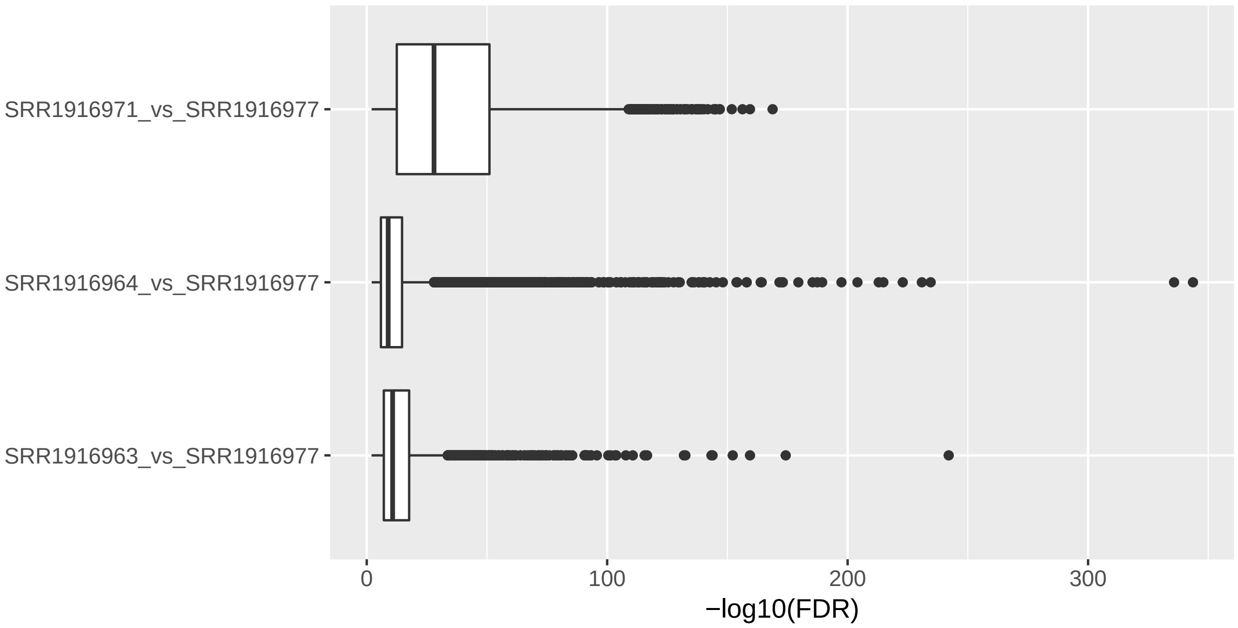


Figure 12: Boxplots of q values for the peaks detected

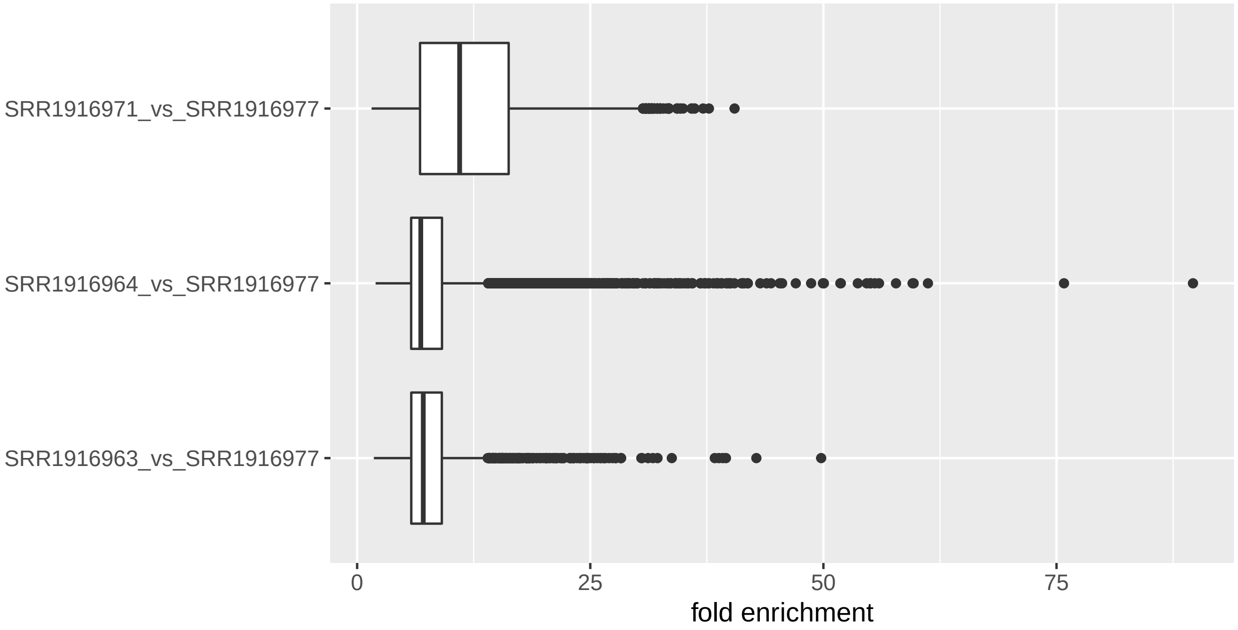


Figure 13: Boxplots of fold change values of peaks detected

##### 3.2.2 Library clustering by similarity (BED mode)

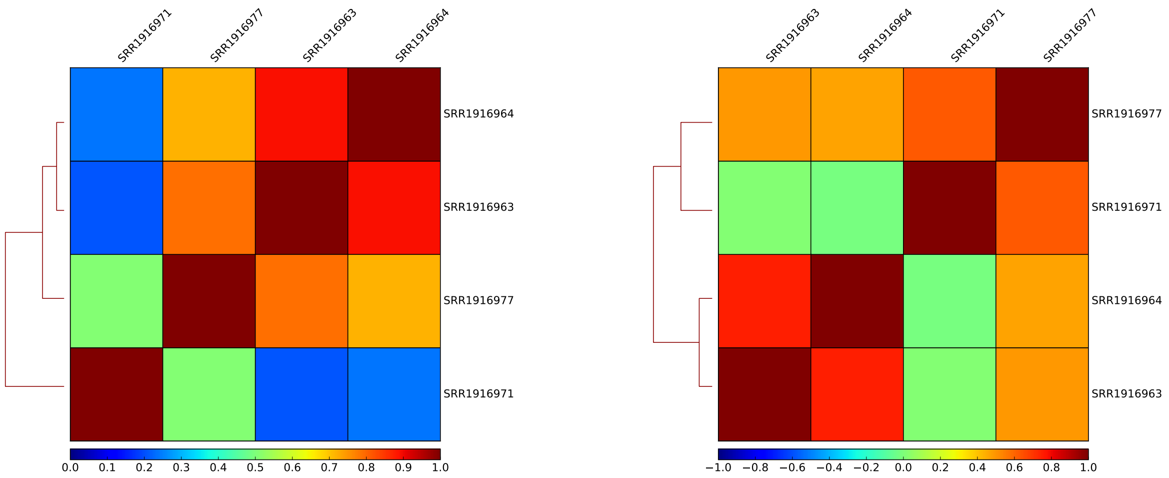


Figure 14: Pearson (left) and Spearman (right) correlation of signal in all libraries, in BED mode, based on the identified peaks regions.

### 4 Results: dataset comparisons

Based on the S1 and S3 libraries a final list peaks was prepared for Pax6, H3K4me3 and SMCX wild type. The final number of peaks, pooled across replicates where applicable is shown in Table 8

| library | peaks |
|---|---|
| H3K4me3_Pax6_SMCX_662.annotated.bed | 922 |
| H3K4me3_Pax6_SMCX_662.annotated.bed.genes.txt | 780 |
| H3K4me3 | 21714 |
| H3K4me3.genes.txt | 18590 |
| Pax6_SMCX_177.annotated.bed.genes.txt | 99 |
| Pax6_SMCX_177.annotated.bed.txt | 194 |
| Pax6_SMCX_662_plus_177.annotated.bed | 874 |
| Pax6_SMCX_662_plus_177.annotated.bed.genes.txt | 675 |
| Pax6 | 10639 |
| Pax6.genes.txt | 6359 |
| SMCX | 10186 |
| SMCX.genes.txt | 7601 |

Table 8: Number of peaks

#### 4.1 Overlapping analyses

##### 4.1.1 Overlapping peaks

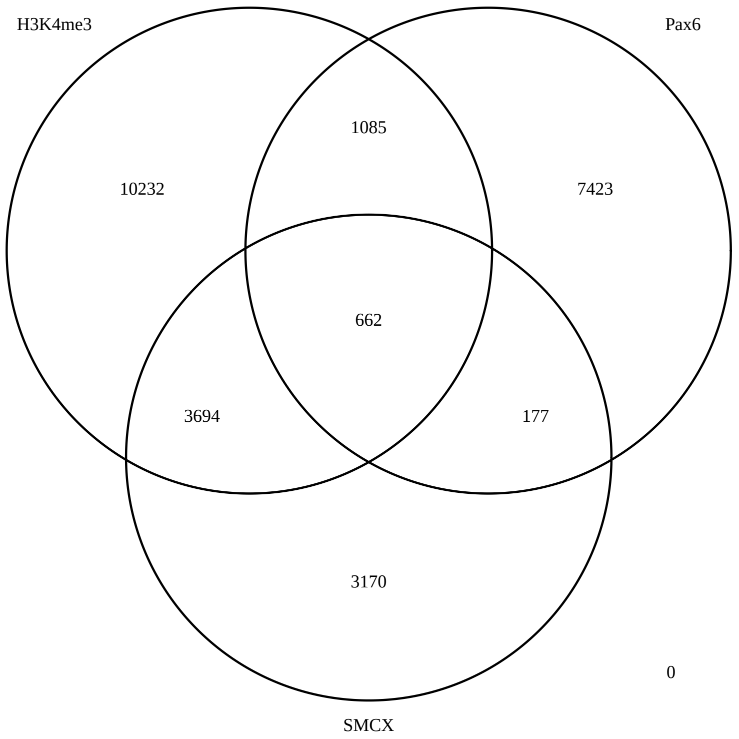


Figure 15: Overlapping peaks between Pax6, H3K4me3 and SMCX

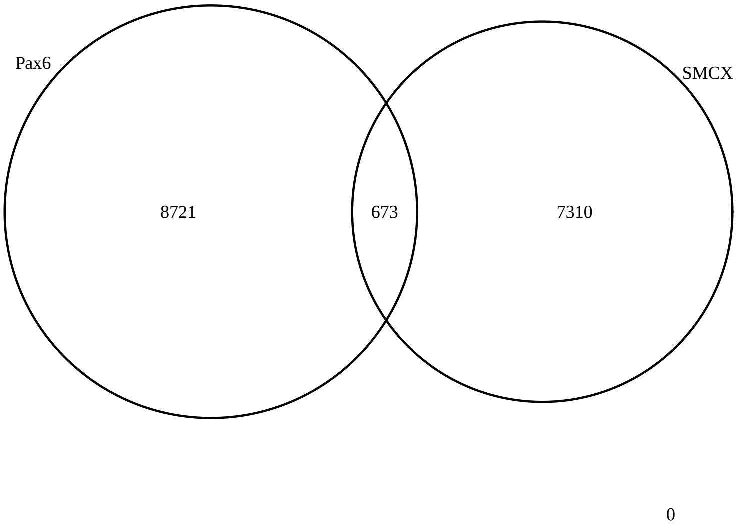


Figure 16: Overlapping peaks between Pax6 and SMCX

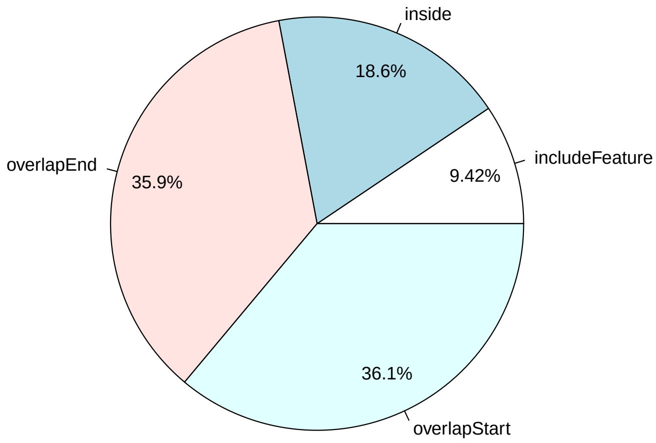


Figure 17: Pie chart of the overlapping features between Pax6 and SMCX

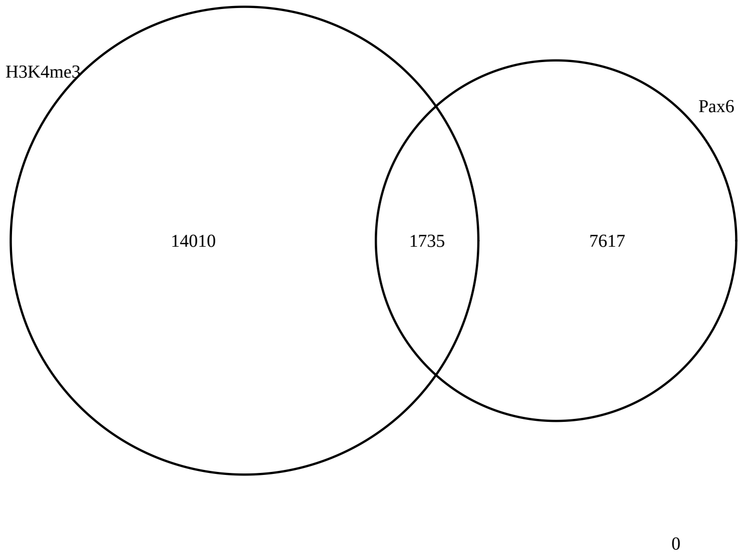


Figure 18: Overlapping peaks between Pax6 and H3K4me3

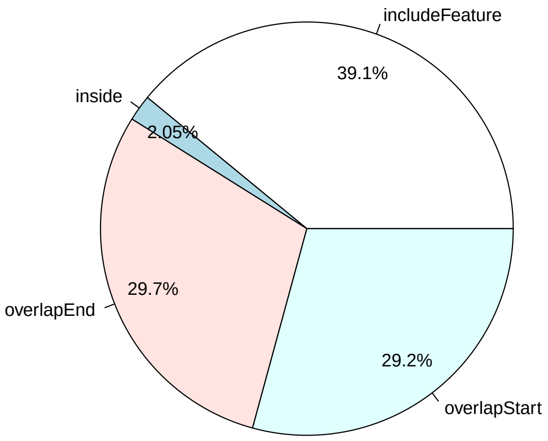


Figure 19: Pie chart of the overlapping features between Pax6 and H3K4me3

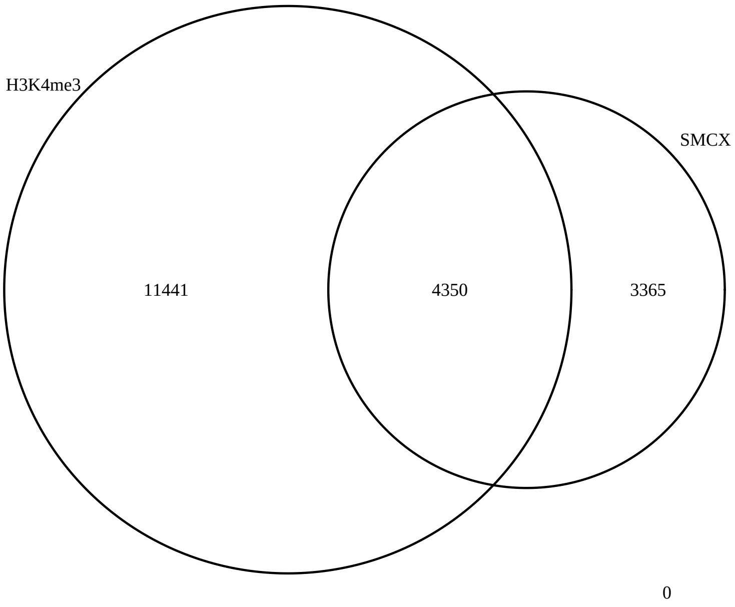


Figure 20: Overlapping peaks between SMCX and H3K4me3

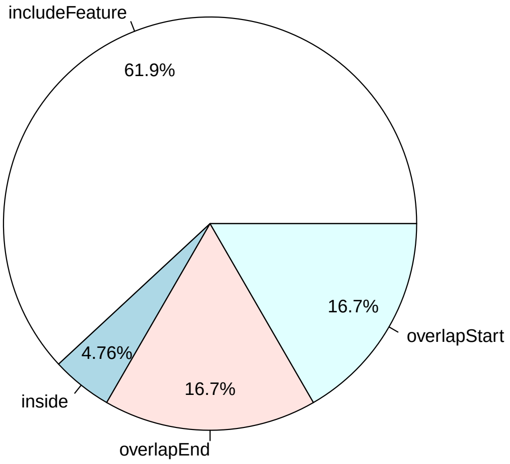


Figure 21: Pie chart of the overlapping features between SMCX and H3K4me3

##### 4.1.2 Overlapping genes

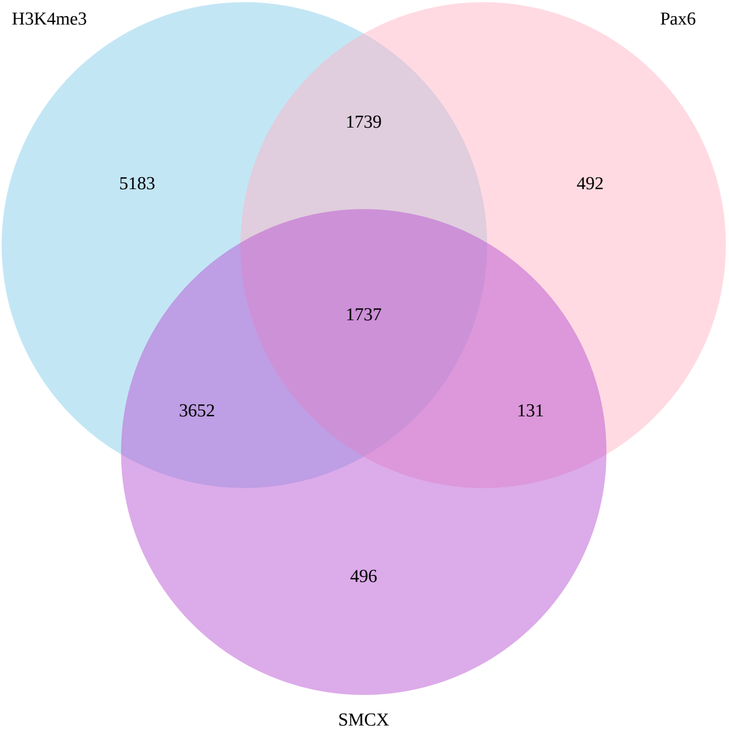


Figure 22: Overlapping genes between Pax6, H3K4me3 and SMCX

#### 4.2 Annotations

| library | GO | PATH |
|---|---|---|
| H3K4me3 | 3326 | 603 |
| Pax6 | 2063 | 92 |
| SMCX | 1257 | 112 |
| Pax6_SMCX_177 | 84 | 4 |
| H3K4me3_Pax6_SMCX_662 | 307 | 2 |
| Pax6_SMCX_662_plus_177 | 267 | 6 |

Table 9: Number of enriched GO and reactome terms given 0.1 BH adjusted p-values

| category | over_represented_pvalue | term | ontology |
|---|---|---|---|
| GO:0005622 | 0.00E+00 | intracellular | CC |
| GO:0005623 | 0.00E+00 | cell | CC |
| GO:0005634 | 0.00E+00 | nucleus | CC |
| GO:0005737 | 0.00E+00 | cytoplasm | CC |
| GO:0043226 | 0.00E+00 | organelle | CC |
| GO:0043227 | 0.00E+00 | membrane-bounded organelle | CC |
| GO:0043229 | 0.00E+00 | intracellular organelle | CC |
| GO:0043231 | 0.00E+00 | intracellular membrane-bounded organelle | CC |
| GO:0044422 | 0.00E+00 | organelle part | CC |
| GO:0044424 | 0.00E+00 | intracellular part | CC |
| GO:0044446 | 0.00E+00 | intracellular organelle part | CC |
| GO:0044464 | 0.00E+00 | cell part | CC |
| GO:0070013 | 4.93E-317 | intracellular organelle lumen | CC |
| GO:0044428 | 1.40E-315 | nuclear part | CC |
| GO:0031974 | 3.71E-315 | membrane-enclosed lumen | CC |

Table 10: Top GO terms, biological processes, for H3K4me3

| category | over_represented_pvalue | term | ontology |
|---|---|---|---|
| GO:0005622 | 0.00E+00 | intracellular | CC |
| GO:0005623 | 0.00E+00 | cell | CC |
| GO:0005634 | 0.00E+00 | nucleus | CC |
| GO:0005737 | 0.00E+00 | cytoplasm | CC |
| GO:0043226 | 0.00E+00 | organelle | CC |
| GO:0043227 | 0.00E+00 | membrane-bounded organelle | CC |
| GO:0043229 | 0.00E+00 | intracellular organelle | CC |
| GO:0043231 | 0.00E+00 | intracellular membrane-bounded organelle | CC |
| GO:0044422 | 0.00E+00 | organelle part | CC |
| GO:0044424 | 0.00E+00 | intracellular part | CC |
| GO:0044446 | 0.00E+00 | intracellular organelle part | CC |
| GO:0044464 | 0.00E+00 | cell part | CC |
| GO:0070013 | 4.93E-317 | intracellular organelle lumen | CC |
| GO:0044428 | 1.40E-315 | nuclear part | CC |
| GO:0031974 | 3.71E-315 | membrane-enclosed lumen | CC |

Table 11: Top GO terms for Pax6

| category | over_represented_pvalue | term | ontology |
|---|---|---|---|
| GO:0005622 | 6.83E-132 | intracellular | CC |
| GO:0044424 | 3.30E-128 | intracellular part | CC |
| GO:0043229 | 6.16E-103 | intracellular organelle | CC |
| GO:0043226 | 1.35E-102 | organelle | CC |
| GO:0044464 | 7.79E-97 | cell part | CC |
| GO:0005623 | 1.13E-96 | cell | CC |
| GO:0005737 | 6.52E-94 | cytoplasm | CC |
| GO:0043227 | 2.44E-88 | membrane-bounded organelle | CC |
| GO:0043231 | 2.14E-86 | intracellular membrane-bounded organelle | CC |
| GO:0044446 | 1.20E-58 | intracellular organelle part | CC |
| GO:0044422 | 3.50E-58 | organelle part | CC |
| GO:0044237 | 3.67E-55 | cellular metabolic process | BP |
| GO:0005488 | 3.39E-53 | binding | MF |
| GO:0044444 | 8.75E-51 | cytoplasmic part | CC |
| GO:0008152 | 2.53E-45 | metabolic process | BP |

Table 12: Top GO terms for SMCX

| category | over_represented_pvalue | term | ontology |
|---|---|---|---|
| GO:0034641 | 1.95E-07 | cellular nitrogen compound metabolic process | BP |
| GO:0031326 | 1.12E-06 | regulation of cellular biosynthetic process | BP |
| GO:0006807 | 1.16E-06 | nitrogen compound metabolic process | BP |
| GO:0019219 | 1.30E-06 | regulation of nucleobase-containing compound metabolic process | BP |
| GO:0009889 | 1.64E-06 | regulation of biosynthetic process | BP |
| GO:0051171 | 1.79E-06 | regulation of nitrogen compound metabolic process | BP |
| GO:0046483 | 2.77E-06 | heterocycle metabolic process | BP |
| GO:1901360 | 3.25E-06 | organic cyclic compound metabolic process | BP |
| GO:0048519 | 3.29E-06 | negative regulation of biological process | BP |
| GO:0006725 | 3.90E-06 | cellular aromatic compound metabolic process | BP |
| GO:0034654 | 4.08E-06 | nucleobase-containing compound biosynthetic process | BP |
| GO:0006139 | 4.67E-06 | nucleobase-containing compound metabolic process | BP |
| GO:0044271 | 5.52E-06 | cellular nitrogen compound biosynthetic process | BP |
| GO:0018130 | 5.71E-06 | heterocycle biosynthetic process | BP |
| GO:0019438 | 6.26E-06 | aromatic compound biosynthetic process | BP |

Table 13: Top GO terms for Pax6 and SMCX (n=177)

| category | over_represented_pvalue | term | ontology |
|---|---|---|---|
| GO:0005622 | 1.50E-30 | intracellular | CC |
| GO:0043231 | 3.57E-30 | intracellular membrane-bounded organelle | CC |
| GO:0044424 | 3.85E-30 | intracellular part | CC |
| GO:0043229 | 4.79E-27 | intracellular organelle | CC |
| GO:0043227 | 4.01E-23 | membrane-bounded organelle | CC |
| GO:0043226 | 1.71E-22 | organelle | CC |
| GO:0070013 | 1.22E-21 | intracellular organelle lumen | CC |
| GO:0031974 | 1.48E-21 | membrane-enclosed lumen | CC |
| GO:0043233 | 1.48E-21 | organelle lumen | CC |
| GO:0044260 | 2.74E-21 | cellular macromolecule metabolic process | BP |
| GO:0005634 | 8.40E-21 | nucleus | CC |
| GO:0043170 | 2.59E-19 | macromolecule metabolic process | BP |
| GO:0044464 | 3.00E-19 | cell part | CC |
| GO:0005623 | 3.15E-19 | cell | CC |
| GO:0044446 | 1.09E-18 | intracellular organelle part | CC |

Table 14: Top GO terms for H3K4me3 and Pax6 and SMCX (n=662)

| category | over_represented_pvalue | term | ontology |
|---|---|---|---|
| GO:0043231 | 5.17E-22 | intracellular membrane-bounded organelle | CC |
| GO:0005622 | 2.93E-21 | intracellular | CC |
| GO:0044424 | 9.01E-21 | intracellular part | CC |
| GO:0034641 | 5.77E-20 | cellular nitrogen compound metabolic process | BP |
| GO:0043229 | 7.40E-19 | intracellular organelle | CC |
| GO:0044260 | 8.64E-19 | cellular macromolecule metabolic process | BP |
| GO:0046483 | 6.82E-18 | heterocycle metabolic process | BP |
| GO:0005634 | 1.18E-17 | nucleus | CC |
| GO:0043227 | 1.33E-17 | membrane-bounded organelle | CC |
| GO:0044237 | 1.58E-17 | cellular metabolic process | BP |
| GO:0006139 | 2.25E-17 | nucleobase-containing compound metabolic process | BP |
| GO:0006725 | 3.58E-17 | cellular aromatic compound metabolic process | BP |
| GO:0006807 | 4.69E-17 | nitrogen compound metabolic process | BP |
| GO:1901360 | 5.61E-17 | organic cyclic compound metabolic process | BP |
| GO:0090304 | 6.63E-17 | nucleic acid metabolic process | BP |

Table 15: Top GO terms for H3K4me3 and Pax6 and SMCX (n=662 plus 177)

| path_name | over_represented_pvalue |
|---|---|
| Mus musculus: Cell Cycle, Mitotic | 4.16E-54 |
| Mus musculus: Gene Expression | 4.97E-53 |
| Mus musculus: Cell Cycle | 1.03E-48 |
| Mus musculus: Organelle biogenesis and maintenance | 5.17E-48 |
| Mus musculus: Assembly of the primary cilium | 1.78E-29 |
| Mus musculus: M Phase | 1.44E-25 |
| Mus musculus: Mitotic Metaphase and Anaphase | 1.42E-23 |
| Mus musculus: Mitotic Anaphase | 2.31E-23 |
| Mus musculus: Separation of Sister Chromatids | 1.09E-21 |
| Mus musculus: Metabolism of proteins | 2.72E-21 |
| Mus musculus: Transcription | 7.58E-21 |
| Mus musculus: Post-translational protein modification | 1.99E-20 |
| Mus musculus: Anchoring of the basal body to the plasma membrane | 5.53E-20 |
| Mus musculus: S Phase | 3.38E-19 |
| Mus musculus: Mitotic G1-G1/S phases | 5.00E-19 |

Table 16: Top reactome terms for H3K4me3

| path_name | over_represented_pvalue |
|---|---|
| Mus musculus: Developmental Biology | 2.17E-13 |
| Mus musculus: Axon guidance | 1.97E-12 |
| Mus musculus: Signaling by NOTCH | 4.01E-08 |
| Mus musculus: Signaling by NOTCH1 | 1.34E-07 |
| Mus musculus: NOTCH1 Intracellular Domain Regulates Transcription | 1.79E-06 |
| Mus musculus: Chromatin modifying enzymes | 7.23E-06 |
| Mus musculus: Chromatin organization | 7.23E-06 |
| Mus musculus: Cell-Cell communication | 7.90E-06 |
| Mus musculus: L1CAM interactions | 1.06E-05 |
| Mus musculus: Cell-cell junction organization | 1.39E-05 |
| Mus musculus: deactivation of the beta-catenin transactivating complex | 1.46E-05 |
| Mus musculus: Fcgamma receptor (FCGR) dependent phagocytosis | 2.02E-05 |
| Mus musculus: Signalling by NGF | 3.01E-05 |
| Mus musculus: Signaling by EGFR | 3.61E-05 |
| Mus musculus: Adherens junctions interactions | 4.21E-05 |

Table 17: Top reactome terms for Pax6

| path_name | over_represented_pvalue |
|---|---|
| Mus musculus: Neuronal System | 3.25E-12 |
| Mus musculus: Organelle biogenesis and maintenance | 4.60E-08 |
| Mus musculus: Voltage gated Potassium channels | 5.69E-08 |
| Mus musculus: Assembly of the primary cilium | 8.27E-08 |
| Mus musculus: Membrane Trafficking | 1.15E-07 |
| Mus musculus: Potassium Channels | 3.42E-07 |
| Mus musculus: Transmission across Chemical Synapses | 4.42E-07 |
| Mus musculus: Axon guidance | 9.07E-07 |
| Mus musculus: Metabolism of nucleotides | 1.25E-06 |
| Mus musculus: Asparagine N-linked glycosylation | 6.72E-06 |
| Mus musculus: Mitotic G2-G2/M phases | 6.94E-06 |
| Mus musculus: Developmental Biology | 1.26E-05 |
| Mus musculus: Fatty acid, triacylglycerol, and ketone body metabolism | 1.62E-05 |
| Mus musculus: G-protein mediated events | 1.94E-05 |
| Mus musculus: G2/M Transition | 2.22E-05 |

Table 18: Top reactome terms for SMCX

| path_name | over_represented_pvalue |
|---|---|
| Mus musculus: Signaling by NOTCH1 | 4.68E-06 |
| Mus musculus: Signaling by NOTCH | 1.26E-05 |
| Mus musculus: NOTCH1 Intracellular Domain Regulates Transcription | 1.74E-05 |
| Mus musculus: Signaling by NOTCH2 | 2.03E-04 |

Table 19: Top reactome terms for Pax6 and SMCX (n=177)

| path_name | over_represented_pvalue |
|---|---|
| Mus musculus: Gene Expression | 1.25E-05 |
| Mus musculus: SMAD2/SMAD3:SMAD4 heterotrimer regulates transcription | 1.03E-04 |

Table 20: Top reactome terms for H3K4me3 and Pax6 and SMCX (n=662)

| path_name | over_represented_pvalue |
|---|---|
| Mus musculus: SMAD2/SMAD3:SMAD4 heterotrimer regulates transcription | 4.62E-05 |
| Mus musculus: Signaling by NOTCH1 | 5.93E-05 |
| Mus musculus: NOTCH1 Intracellular Domain Regulates Transcription | 9.83E-05 |
| Mus musculus: Signaling by NOTCH | 2.86E-04 |
| Mus musculus: Fcgamma receptor (FCGR) dependent phagocytosis | 2.86E-04 |
| Mus musculus: Membrane Trafficking | 3.11E-04 |

Table 21: Top reactome terms for H3K4me3 and Pax6 and SMCX (n=662 plus 177)

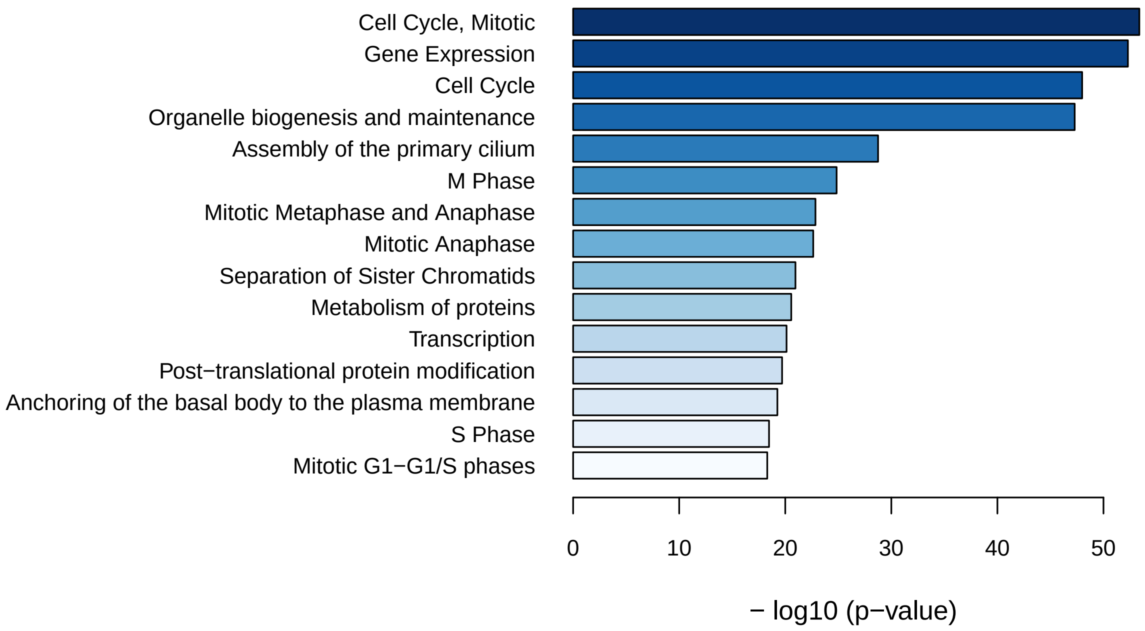


Figure 23: Top reactome terms for H3K4me3

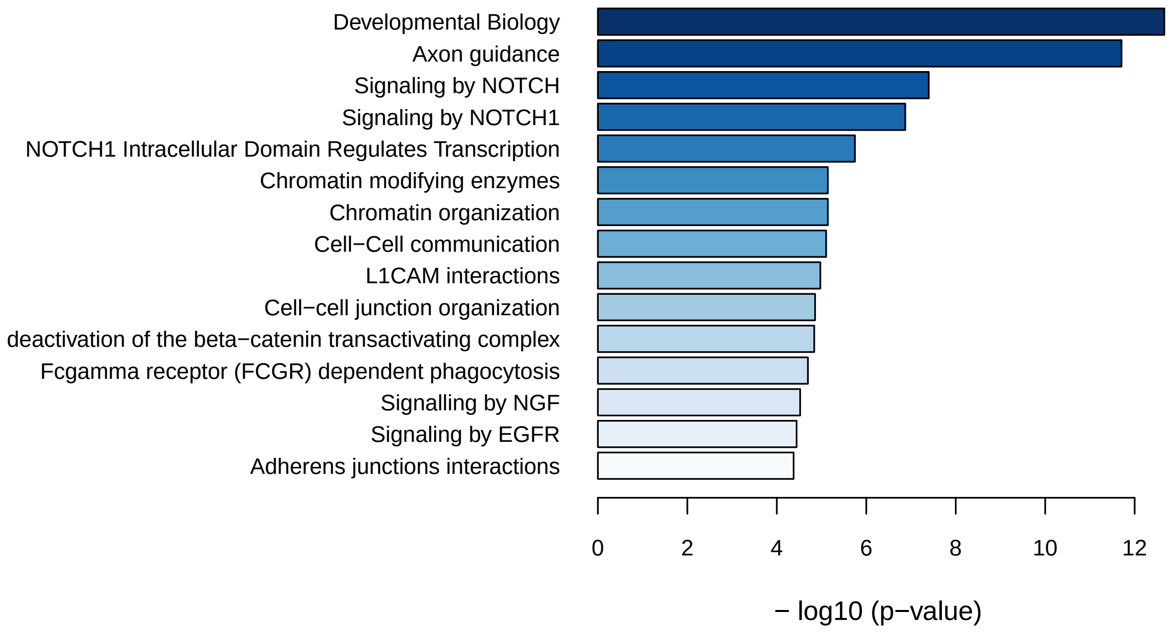


Figure 24: Top reactome terms for Pax6

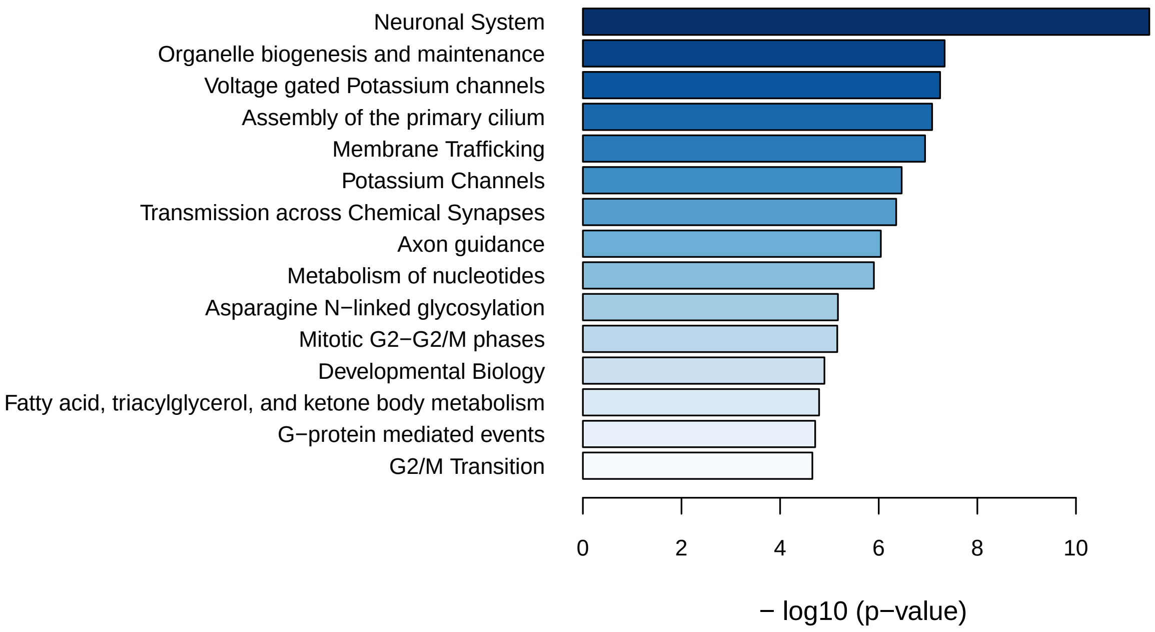


Figure 25: Top reactome terms for SMCX

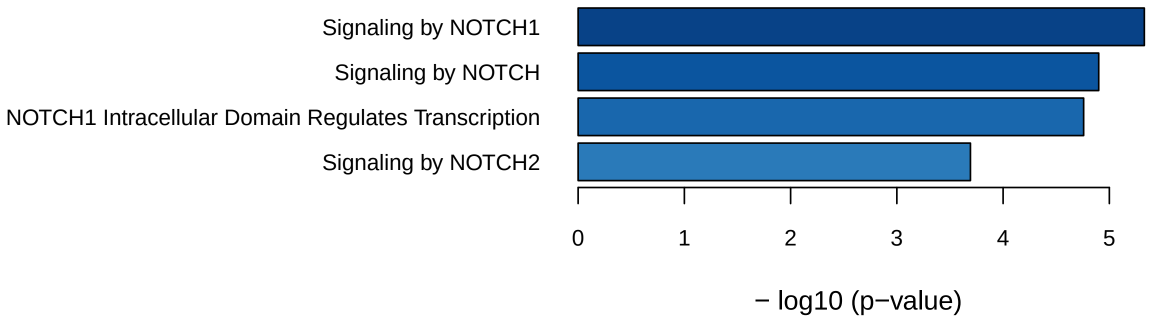


Figure 26: Top reactome terms for Pax6_SMCX_177

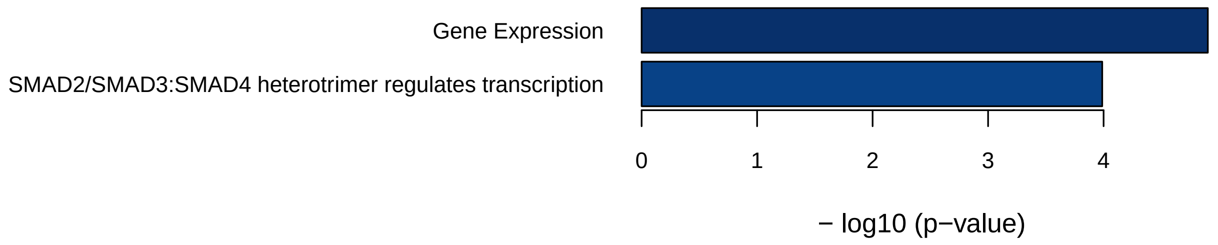


Figure 27: Top reactome terms for H3K4me3_Pax6_SMCX_662

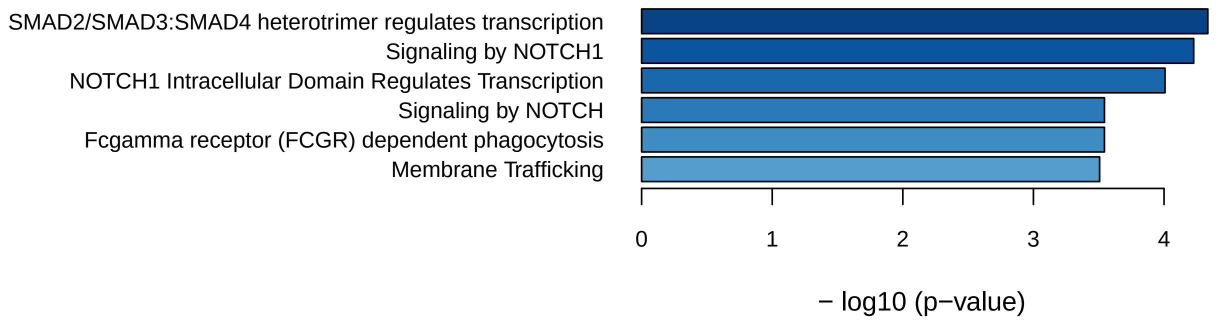


Figure 28: Top reactome terms for Pax6_SMCX_662_plus_1772

##### 4.2.1 Peaks distribution over features

Summarizing peak distribution over exon, intron, enhancer, proximal promoter, 5 prime UTR and 3 prime UTR was done using `assignChromosomeRegion` function in the `ChIPpeakAnno_3.8.9` package.

The counting was done twice:

1. enabling double counting, meaning that if a peak overlaps with both promoter and 5ÚTR, both promoter and 5ÚTR are counted. Here, the overall percentage over features will exceed 100% if any of peaks spams over two regions

2. disabling double counting by specifying a precedence order. For example, if promoter is specified before 5ÚTR, then only promoter will be incremented for the same example. The precedence used was set to default and included promoters, immediateDownstream, fiveUTRs, threeUTRs, exons and introns in this particular order.

In addition, a default cutoff of 1000 bases was used to immediate downstream region. Peaks that reside within immediate downstream cutoff downstream of gene end but not overlap with

3 prime UTR are classified as immediate downstream. Peaks that reside downstream over immediate downstream cutoff from gene end are classified as intergenic, potential enhancers or silencers.

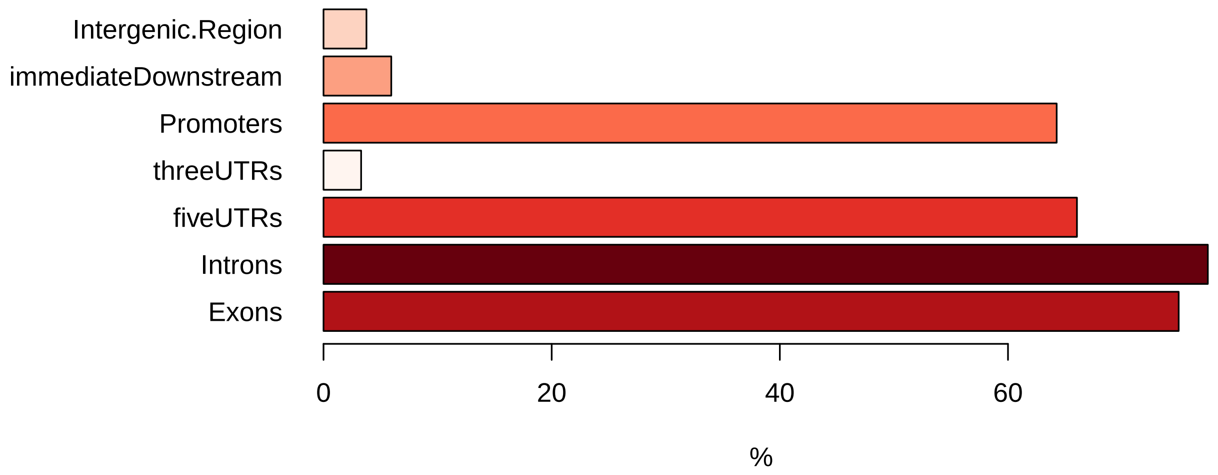


Figure 29: Peaks distribution over features for H3K4me3, double count are enabled for peaks overlapping more than one feature (e.g. promoter and 5'UTR)

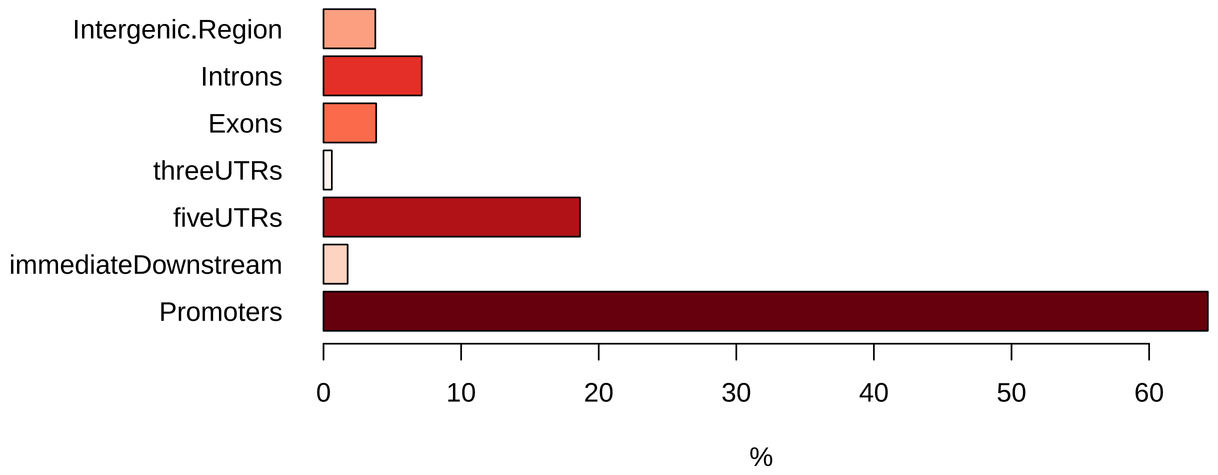


Figure 30: Peaks distribution over features for H3K4me3, counts precedence: promoters, immediateDownstream, fiveUTRs, threeUTRs, exons and introns)

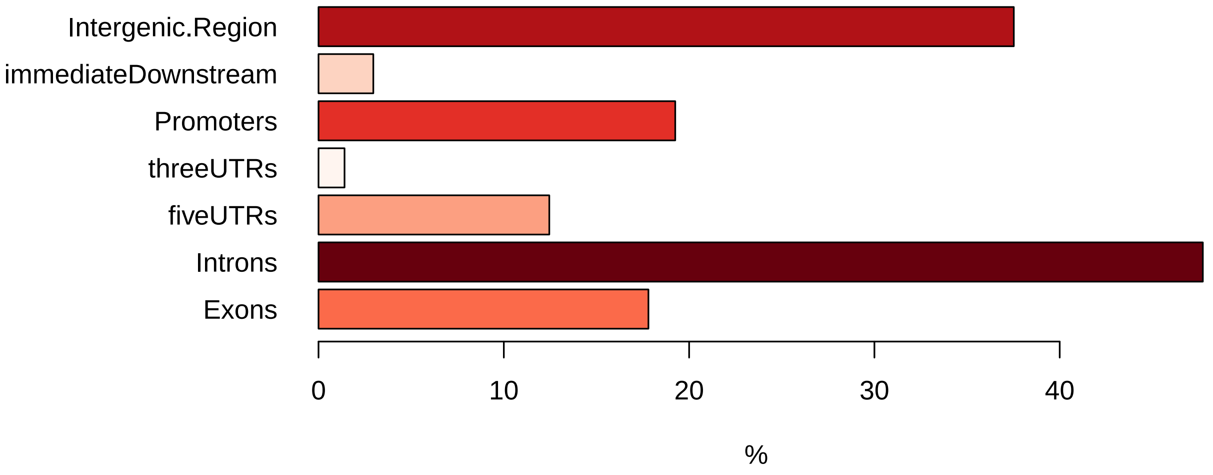


Figure 31: Peaks distribution over features for Pax6, double count are enabled for peaks overlapping more than one feature (e.g. promoter and 5'UTR)

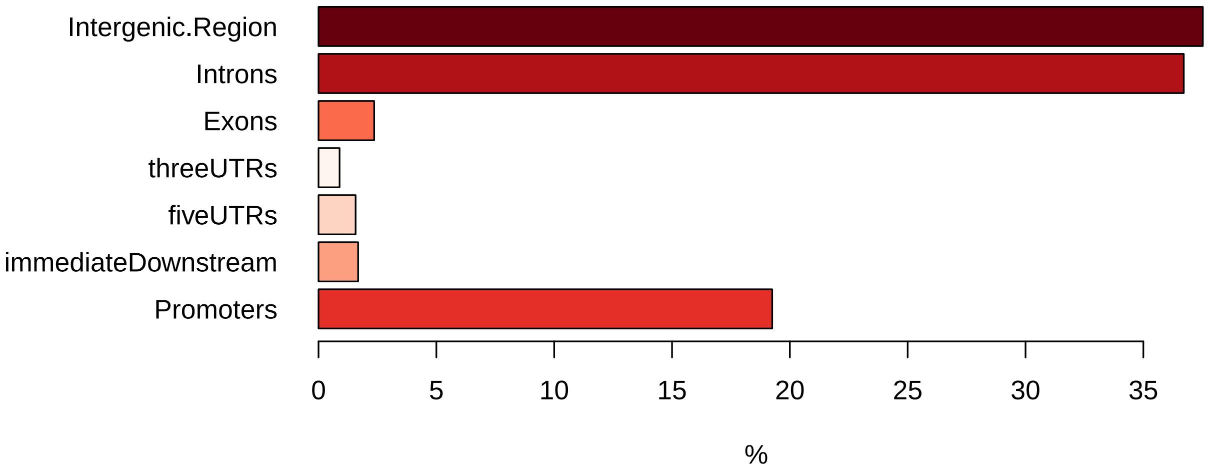


Figure 32: Peaks distribution over features for Pax6, counts precedence: promoters, immediateDownstream, fiveUTRs, threeUTRs, exons and introns)

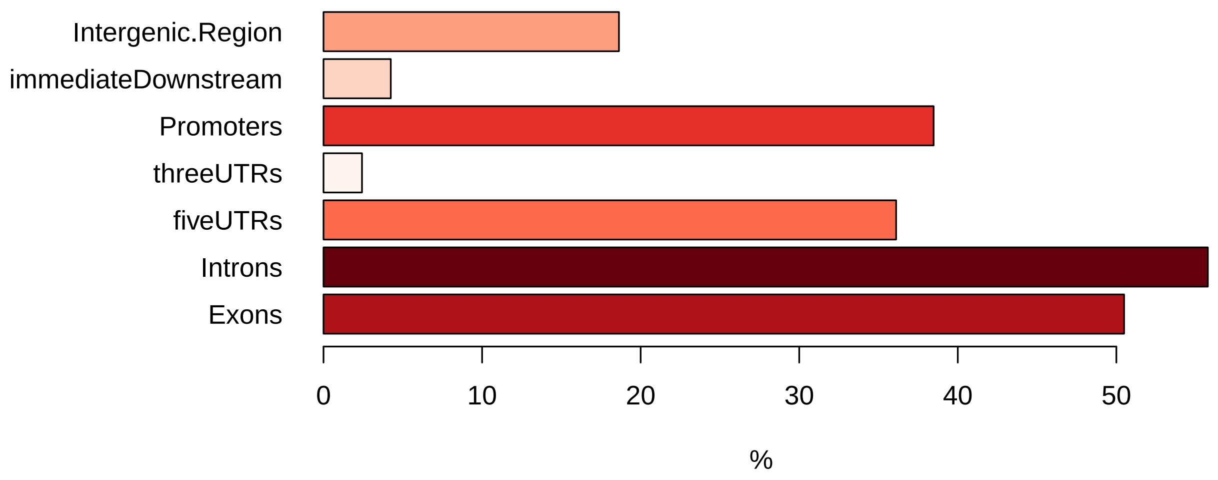


Figure 33: Peaks distribution over features for SMCX, double count are enabled for peaks overlapping more than one feature (e.g. promoter and 5'UTR)

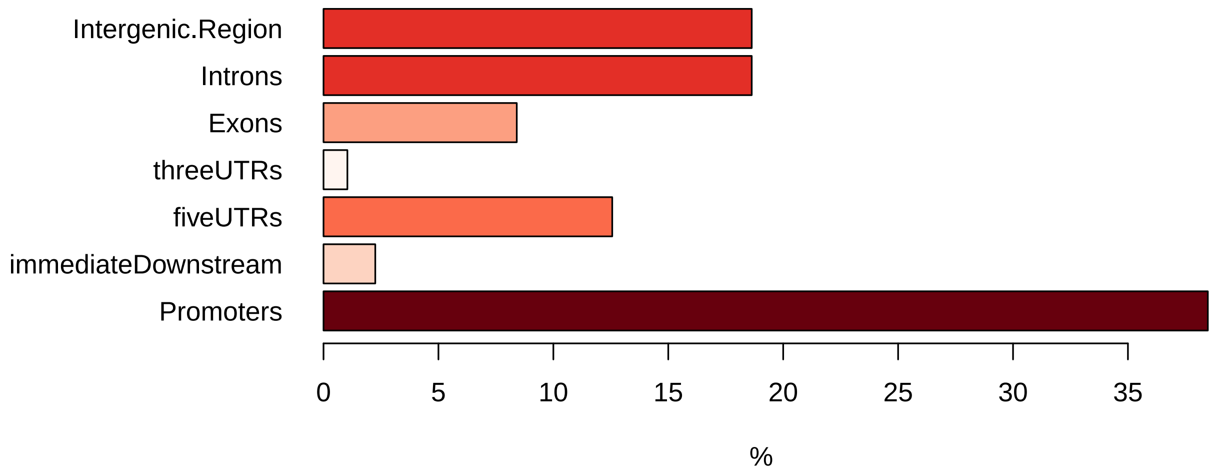


Figure 34: Peaks distribution over features for SMCX, counts precedence: promoters, immediateDownstream, fiveUTRs, threeUTRs, exons and introns)

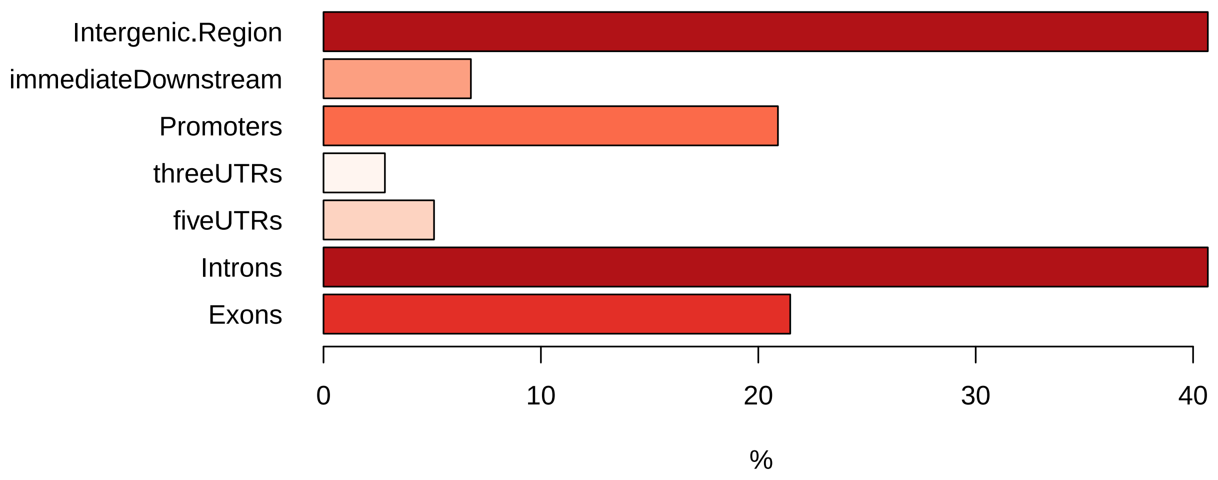


Figure 35: Peaks distribution over features for Pax6_SMCX_177, double count are enabled for peaks overlapping more than one feature (e.g. promoter and 5'UTR)

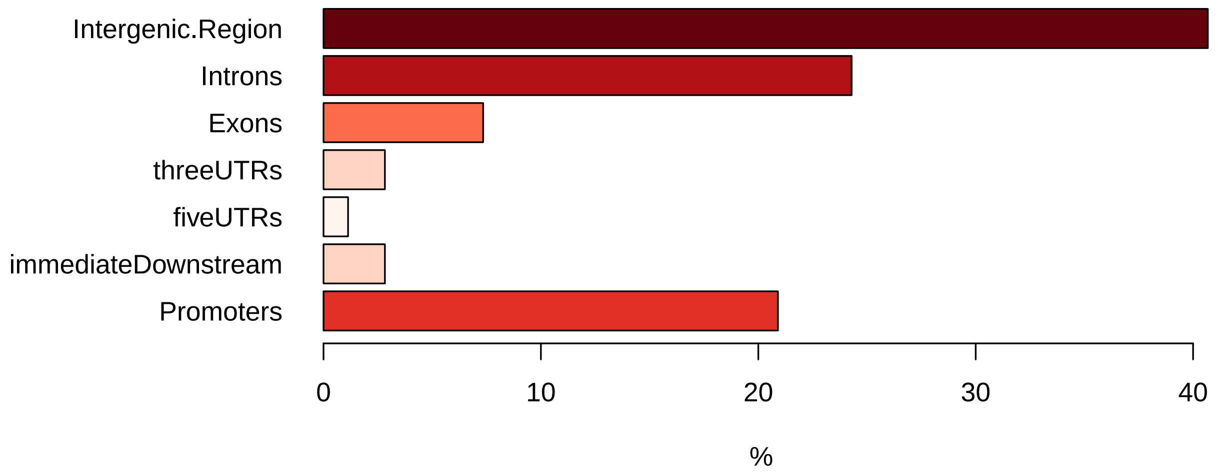


Figure 36: Peaks distribution over features for Pax6_SMCX_177, counts precedence: promoters, immediateDownstream, fiveUTRs, threeUTRs, exons and introns)

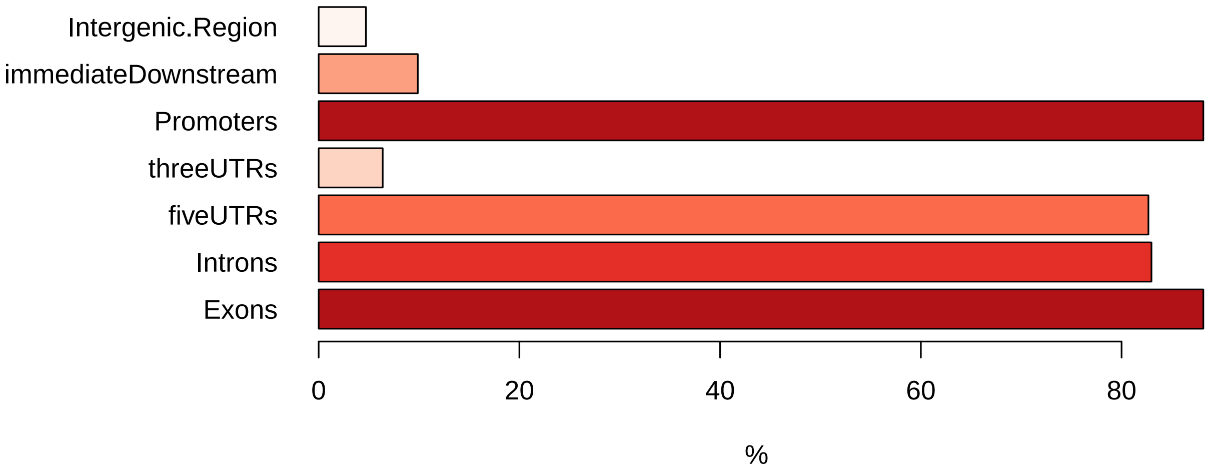


Figure 37: Peaks distribution over features for H3K4me3_Pax6_SMCX_662, double count are enabled for peaks overlapping more than one feature (e.g. promoter and 5'UTR)

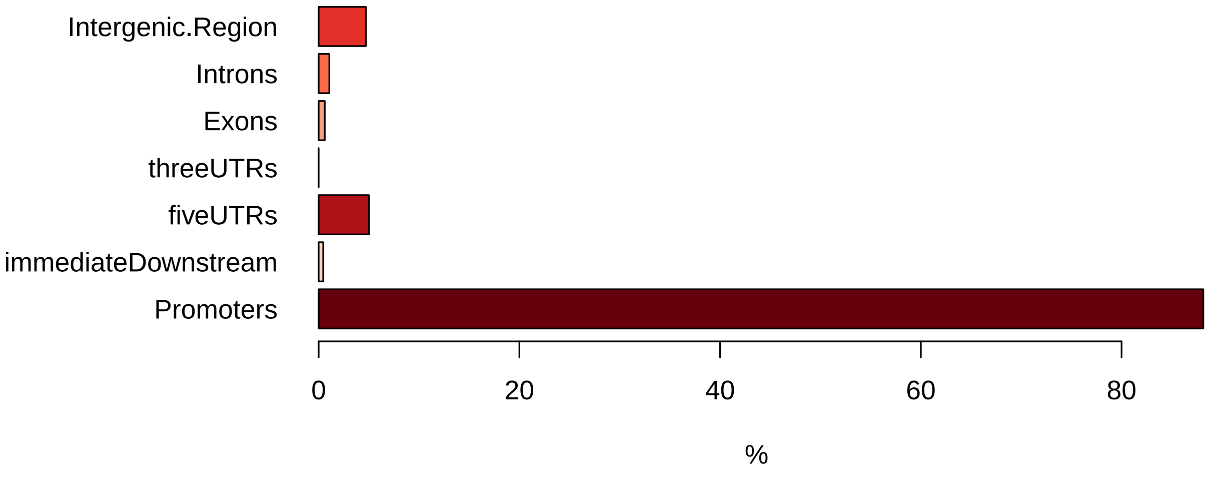


Figure 38: Peaks distribution over features for H3K4me3_Pax6_SMCX_662, counts precedence: promoters, immediateDownstream, fiveUTRs, threeUTRs, exons and introns)

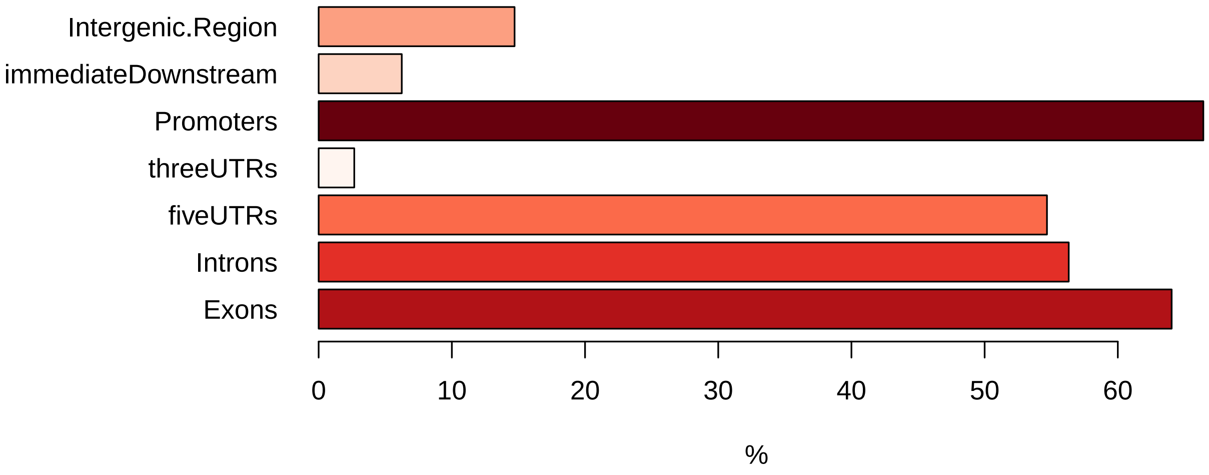


Figure 39: Peaks distribution over features for Pax6_SMCX_662_plus_177, double count are enabled for peaks overlapping more than one feature (e.g. promoter and 5'UTR)

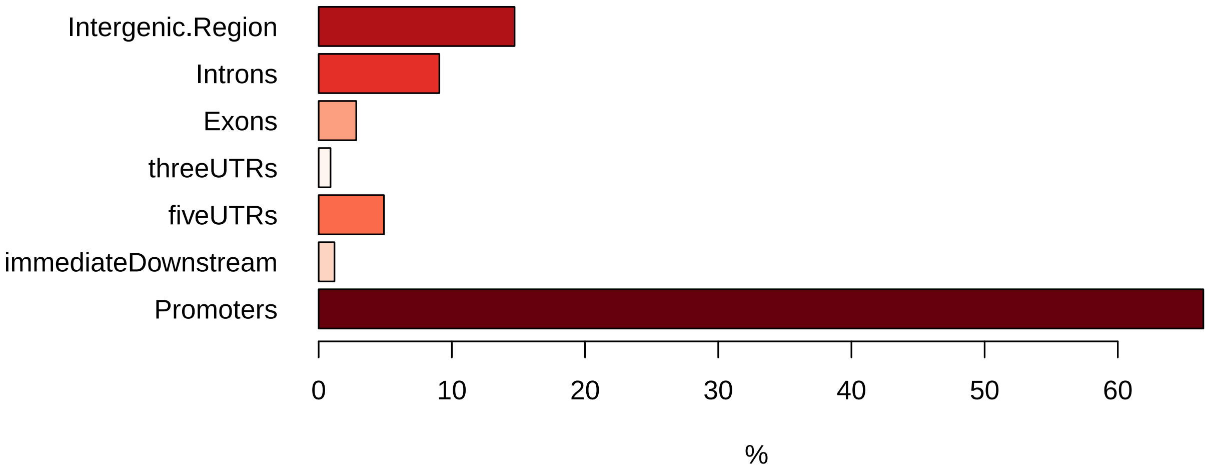


Figure 40: Peaks distribution over features for Pax6_SMCX_662_plus_177, counts precedence: promoters, immediateDownstream, fiveUTRs, threeUTRs, exons and introns)

BroadInstitute. *Picard*. https://broadinstitute.github.io/picard/.

Phantompeakqualtools. *Phantompeakqualtools*. https://www.encodeproject.org/software/phantompeak.

Ramírez, Fidel, et al. 2014. "DeepTools: A flexible platform for exploring deep-sequencing data". *Nucleic Acids Research* 42 (W1): 187–191.

Feng, Jianxing, et al. 2012. "Identifying ChiP-seq enrichment using MACS". *Nat. Protocols* 7 (9): 1728–1740.

Neph, Shane, et al. 2012. "BEDOPS: High-performance genomic feature operations". *Bioinformatics* 28 (14): 1919–1920.

Zhu, Lihua J, et al. 2010. "ChIPpeakAnno: a Bioconductor package to annotate ChIP-seq and ChIP-chip data". *BMC Bioinformatics* 11 (1): 237. ISSN: 1471-2105. doi:10.1186/1471-2105-11-237. http://dx.doi.org/10.1186/1471-2105-11-237.

Young, Matthew D, et al. 2010. "Gene ontology analysis for RNA-seq: accounting for selection bias." *Genome biology* 11 (2): R14. ISSN: 1465-6914. doi:10.1186/gb-2010-11-2-r14. http://www.pubmedcentral.nih.gov/articlerender.fcgi?artid=2872874{\&}tool=pmcentrez{\&}rendertype=abstract.

### R session info

- R version 3.3.3 (2017-03-06), `x86_64-apple-darwin13.4.0`
- Base packages: base, datasets, grDevices, graphics, methods, stats, utils
- Other packages: RColorBrewer 1.1-2, ggplot2 2.2.1, knitr 1.17, xtable 1.8-2
- Loaded via a namespace (and not attached): Rcpp 0.12.13, colorspace 1.3-2, evaluate 0.10.1, grid 3.3.3, gtable 0.2.0, labeling 0.3, lazyeval 0.2.1, magrittr 1.5, munsell 0.4.3, plyr 1.8.4, rlang 0.1.4, scales 0.5.0, stringi 1.1.6, stringr 1.2.0, tibble 1.3.4, tools 3.3.3
